# Priming Epigenetic Landscape at Gene Promoters through Transcriptional Activation in Mammalian Germ Cells

**DOI:** 10.1101/2024.11.24.625111

**Authors:** Peilin Li, Tatsuya Fujisawa, Haruka Narita, Yuta Uneme, Masahide Seki, Mengwen Hu, Ryohei Narumi, Satoshi H. Namekawa, Shehroz S. Khan, James R. Davie, Yutaka Suzuki, Jun Adachi, Ahmed Ashraf, Azusa Inoue, Mikiko C. Siomi, Soichiro Yamanaka

## Abstract

In mammalian germ cells, a cycle of erasing and resetting epigenetic information is crucial for sex-specific gametogenesis and early embryogenesis. Specific chromatin states at promoters allow genes to be expressed at the appropriate time during germ cell development, however, the mechanism that establishes such characteristic chromatin states remains unclear. Here, we show that, in mouse male embryonic germ cells, promoters of developmental and housekeeping genes are selectively protected from *de novo* DNA methylation, coinciding with their transient upregulation and genome-wide erasure of H3K27me3. Importantly, a specific level of H3K4me3 density at the promoters serves as a potent probe that distinguishes between hypo- and hyper-DNA methylation states. Subsequent restoration of H3K27me3 contributes to establishing bivalent chromatin states that prime genes for activation at later developmental stages. These findings reveal a molecular framework shaping the epigenetic landscape at promoters with long-term effects beyond germ cell development.

## Introduction

Germ cells are the only cell type capable of transmitting genetic and epigenetic information necessary for the development of individuals across generations. During gametogenesis, governing appropriate gene expression is critical for ensuring reproductive competence. In the mammalian germline, gene expression is largely regulated by two types of epigenetic marks at gene loci: DNA methylation and histone modifications. While DNA methylation stably represses gene transcription, histone modifications flexibly switch gene expression between “ON” and “OFF” states according to developmental stages. This indicates that the two epigenetic marks constitute different layers of gene regulation, though their functional relationships in germ cells remain unclear.

DNA methylation undergoes drastic reprogramming during gametogenesis in mammals. Following the specification of mouse primordial germ cells (PGCs), DNA methylation is globally erased, resulting in the most hypomethylated state observed throughout the life cycle ^1–4^. In the male germline, genome-wide DNA methylation is subsequently re-established in gonocytes, also known as prospermatogonia, which emerge between embryonic day (E) 13.5 and postnatal day (P) 3, serving as essential intermediates between PGCs and spermatogonia. This process, referred to as *de novo* DNA methylation, is essential for male fertility, as its defects lead to severe hypogonadism ^5^. *De novo* DNA methylation exhibits a lenient preference for gene bodies and intergenic regions ^6–8^. It implies that gene promoter regions may resist such modification in gonocytes, though the underlying mechanisms remain largely unknown.

Compared to the regulatory layer mediated by DNA methylation, the dynamic regulation of histone modifications during development is critical for cellular and tissue-specific gene expression. Spermatogonial stem cells (SSCs), derived from gonocytes, sustain lifelong sperm production through continuous self-renewal and differentiation, a process largely maintained by a delicate balance of histone modifications. Specifically, two different Polycomb repressive complexes (PRCs) play key roles in this regulation. PRC2 (Polycomb repressive complex 2) deposits H3K27me3 on specific gene groups, repressing soma-specific transcription and maintaining germ cell homeostasis ^9,10^. PRC1, which catalyzes H2AK119ub, directs the timely activation of germline genes and mediates H3K27me3 deposition at specific loci to prevent SSCs from differentiating ^11,12^.

Moreover, the co-occupancy of the repressive mark H3K27me3 and the permissive mark H3K4me3 at gene promoters represents the “bivalent state”. This bivalent state is proposed to prime silenced genes for rapid activation upon cell differentiation, thereby contributing to epigenetic priming in various cell lineages, including SSCs ^13–19^. It creates stable yet reversible chromatin states that allow proper spermatogenesis in the male germline ^20^. Though, it remains elusive how the SSC-specific epigenetic primed state is established in male germline cells. Accordingly, we hypothesized that in gonocytes, essential gene promoters may undergo at least two prerequisite steps to achieve coordinated expression in the male germline. The first involves avoiding *de novo* DNA methylation, and the second entails establishing SSC-specific bivalent states at these hypo-methylated promoters. Therefore, we aimed to investigate how these two steps are executed and sequentially coordinated in gonocytes to ensure proper gene expression in the male germline.

In this study, we analyzed DNA methylome data of mouse male embryonic germ cells and found that the majority of gene promoters, unlike gene bodies and intergenic regions, remain hypomethylated throughout the *de novo* DNA methylation. By examining gene expression changes during this process, we unexpectedly observed a transient and ectopic upregulation of thousands of genes during the gonocyte middle stage. Intriguingly, H3K27me3 and H2AK119ub, along with genomic binding level of some Polycomb complex subunits, underwent a global loss during this period, which may underpin the gene upregulation. Furthermore, we implied that the dynamic increase in H3K4me3 density correlates with the *de novo* DNA methylation level at promoters, contributing to the avoidance of both developmental and housekeeping genes from *de novo* DNA methylation. It also indicated that a specific level of H3K4me3 density serves as a robust probe to distinguish between hypo or hyper DNA methylation at promoters. We also illustrated how the SSC-specific chromatin state was established during the gonocyte-to-spermatogonia transition and suggested that the epigenetic reprogramming in gonocytes may prime them for subsequent developmental processes, including spermatogenesis and preimplantation embryogenesis. These findings highlighted that the perinatal chromatin dynamics in male germ cells may have long-term effects on mature sperm and the next generation.

## Results

### Promoters of Highly Expressed Genes Resist *De Novo* DNA Methylation

In the mouse male germline, *de novo* DNA methylation begins at E13.5 and progressively establishes across the genome by P0 ^21^ (Fig. 1A). To investigate how DNA methylation is established in different chromatin contexts, we analyzed publicly available whole-genome bisulfite sequencing (WGBS-seq) data ^6,22^, examining DNA methylation levels at E13.5, E16.5, and P0 in three genomic regions: promoters, gene bodies, and intergenic regions. At E13.5, DNA methylation levels were relatively low across all three regions (Fig. 1B), consistent with the global loss of DNA methylation in PGCs ^1^. From E13.5 to P0, DNA methylation levels drastically increased in gene bodies and intergenic regions (Fig. 1B). In contrast, promoters remained less methylated throughout the period, suggesting the existence of a mechanism that maintains such hypomethylated state at promoters during the gonocyte period (Fig. 1B).

**Fig. 1:**
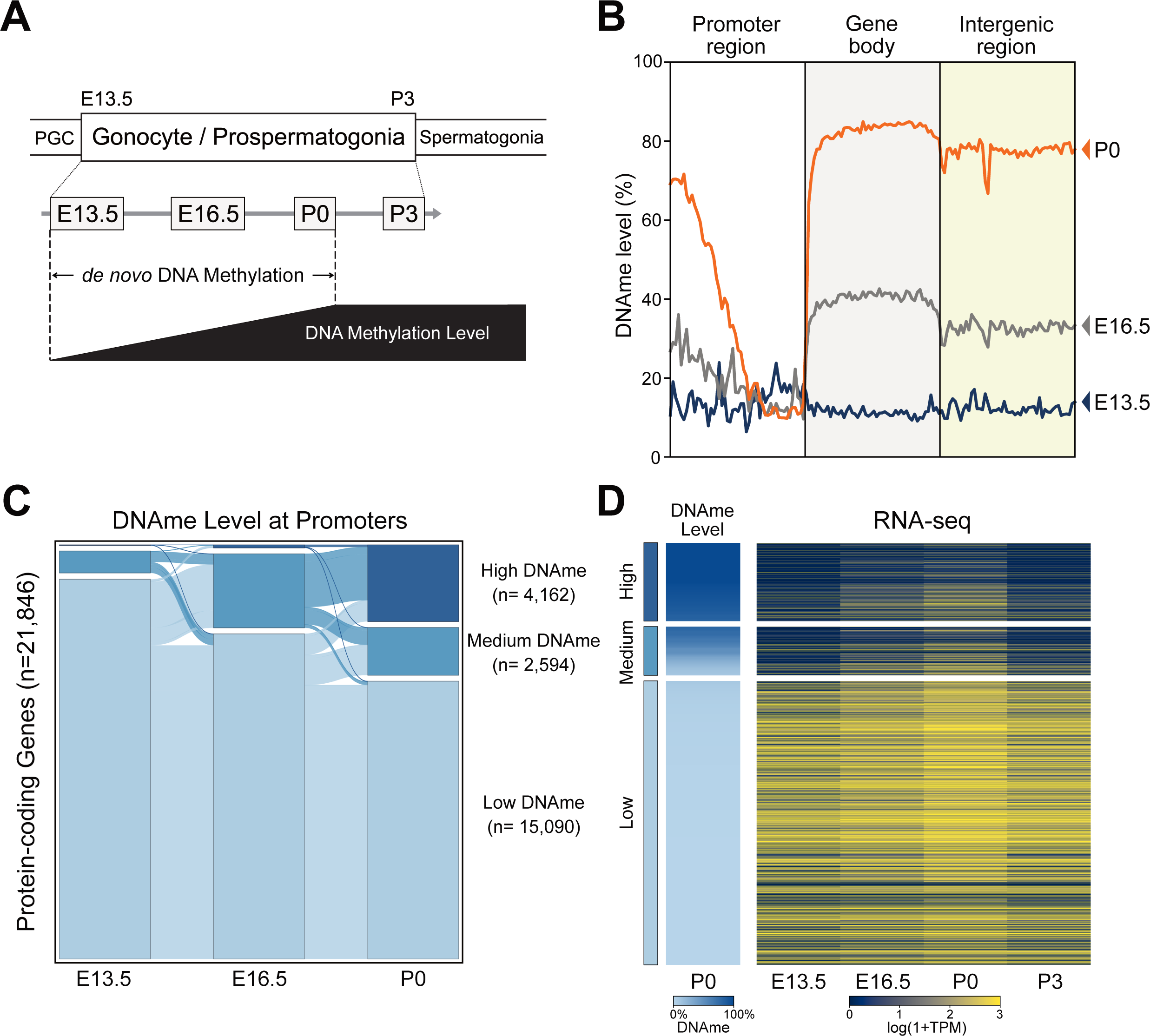
Promoters of Highly Expressed Genes Resist *De Novo* DNA Methylation. (A) Schematic of spermatogenesis and *de novo* DNA methylation. Gonocytes represent embryonic male germ cells from embryonic day (E) 13.5 to postnatal day (P) 3. *De novo* DNA methylation starts at E13.5 and finishes around P0. (B) Line plots displaying average DNA methylation levels from E13.5 to P0 across the mouse genome categorized into either promoter regions (TSS -2000bp to +500bp), gene body regions (gene regions excluding promoters), and intergenic regions (genome regions excluding genes and promoters). (C) Sankey diagram showing changes in DNA methylation levels of promoter regions (TSS +/- 500bp) for all protein-coding genes (n=21,846) from E13.5 to P0. Promoters are categorized into three groups as Low (0% - 10%), Medium (10% - 80%), and High (80% - 100%) DNA methylation (DNAme), respectively, based on DNA methylation level at each stage. (D) Heatmap illustrating the correlation between P0 DNA methylation levels and gene expression levels at each stage. Promoters sorted by P0 DNA methylation levels and categorized as defined in (**C**) are used for analysis. See also Figure S1.

We then analyzed the changes in DNA methylation levels at each promoter of 21,846 protein-coding genes during the gonocyte period, revealing that nearly 70% of promoters avoid *de novo* DNA methylation by P0 (Fig. 1C). Based on the DNA methylation levels at P0, all promoters were categorized into three groups, termed High, Medium, and Low DNA methylation (DNAme) (Fig. 1C). The High and Medium DNAme groups, containing 2,594 (11.9%) and 4,162 promoters (19.1%), respectively, gradually acquired DNA methylation from E13.5 to P0 (Fig. 1C). Gene Ontology (GO) analysis indicated that most of these promoters are linked to genes involved in the immune system or sensory system, such as olfactory receptor genes ^23,24^ and vomeronasal receptor genes ^25^ (Fig. S1A, B). These genes tend to be located near transposable elements (Fig. S1D), which are largely silenced through *de novo* DNA methylation in gonocytes ^26^. On the other hand, genes in the Low DNAme group (n=15,090) are predominantly associated with developmental and housekeeping genes (Fig. 1C, Fig. S1C). This implied that the expression of these genes is likely preserved during *de novo* DNA methylation.

Next, to understand how *de novo* DNA methylation affects gene expression, we examined the changes in expression level of each gene in the three groups defined above using our previous RNA-seq data ^27^ (Fig. 1D). It revealed a trend where genes with lower expression levels were generally more likely to be methylated, while genes in Low DNAme group exhibited higher transcriptional activity, even at E13.5, prior to the onset of *de novo* DNA methylation (Fig. 1D). This implied that the extent to which a gene acquires *de novo* DNA methylation in male germ cells is influenced by factors associated with gene activity.

### Diverse Genes Experience Transient and Ectopic Upregulation in Gonocytes

We noticed that all three groups defined above (High, Medium, and Low DNAme) showed upregulation toward the middle stage of gonocyte (E16.5 - P0) (Fig. 1D). This finding was contrary to our expectation, considering the *de novo* DNA methylation is in progress during this period. To reveal how gene expression is regulated in gonocytes, we compared RNA-seq data between each pair of adjacent stages within the gonocyte stage (E13.5 vs E16.5, E16.5 vs P0, and P0 vs P3) ^27^. Accordingly, 2,407 genes were upregulated from E13.5 to E16.5, and 1,523 genes were upregulated from E16.5 to P0 (Fig. 2A, B), followed by the downregulation of 3,097 genes from P0 to P3 (Fig. 2A, B). Genes upregulated from E13.5 to P0 showed a considerable overlap with those downregulated from P0 to P3 (Fig. 2C, S2A). Moreover, genes whose expression increased between E13.5 and E16.5, underwent downregulation after P0 (Fig. 2D). These findings indicated that thousands of genes are transiently activated during the middle stage of gonocyte.

**Fig. 2:**
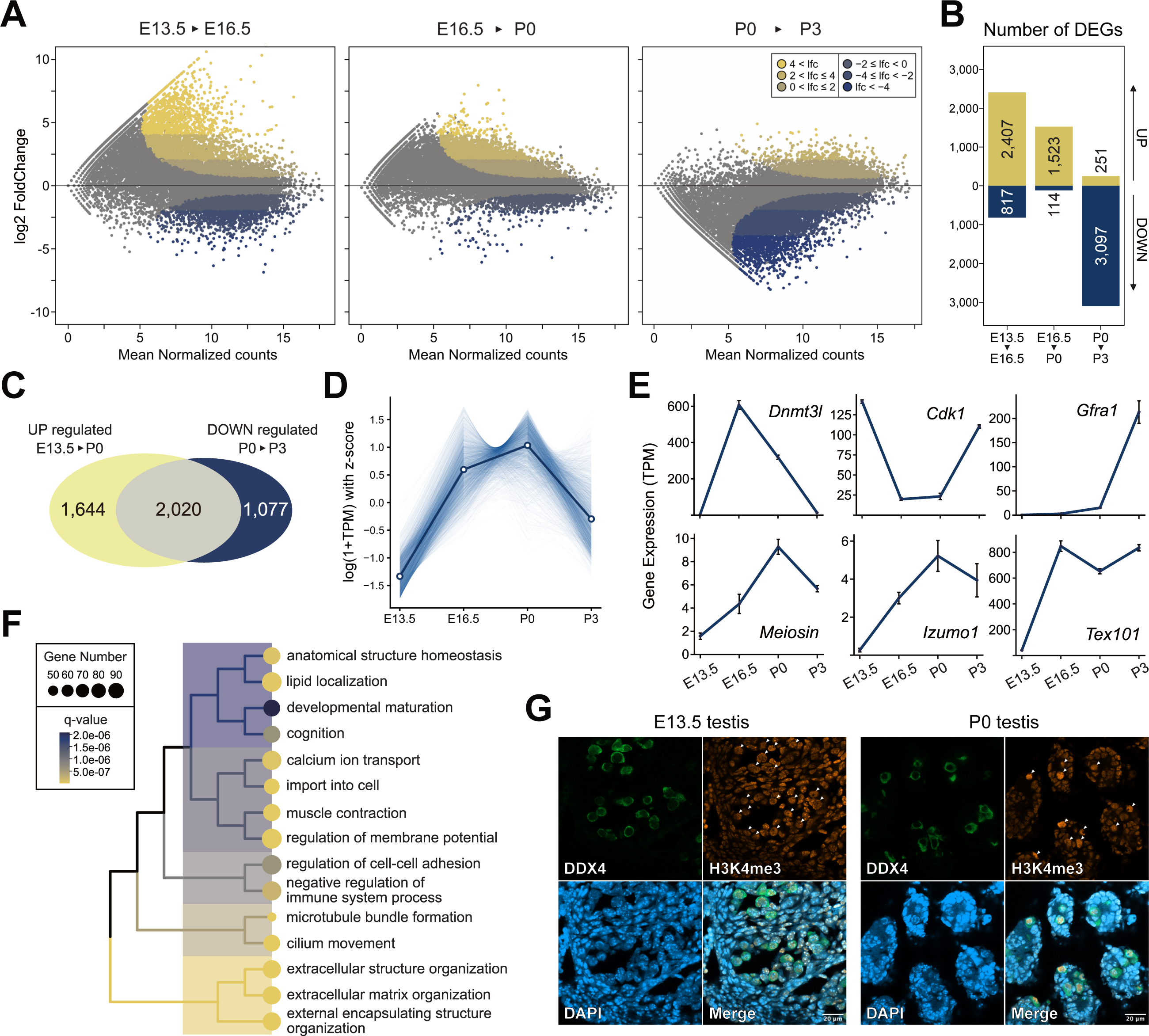
Diverse Genes Experience Transient and Ectopic Upregulation in Gonocytes. (**A**) MA-plots comparing expression of all protein-coding genes between each adjacent stages during gonocyte period: E13.5 to E16.5, E16.5 to P0, and P0 to P3. Genes significantly up- or downregulated (q-value < 1e-12) with varying log2FoldChange thresholds are annotated. (**B**) Number of differentially expressed genes (DEGs) between each adjacent stages. Upregulated genes (yellow) and downregulated genes (blue) are identified by q-value < 1e-12 and log2FoldChange > 2 or < -2, respectively. (**C**) Venn diagram illustrating the overlap between upregulated DEGs from E13.5 to P0 (n=3,664, combining upregulated DEGs from E13.5 to E16.5 and from E16.5 to P0) and downregulated DEGs from P0 to P3 (n=3,097). (**D**) Integrated line plot showing expression changes of upregulated DEGs from E13.5 to E16.5 (n=2,407) during gonocyte period. (**E**) Line plots showing gene expression changes from E13.5 to P3, marker genes (first row) and ectopically expressed genes (second row) are represented. (**F**) Gene Ontology analysis of upregulated DEGs from E13.5 to E16.5 (n=2,407). Top fifteen terms are listed and annotated with hierarchical clusters. (**G**) Seminiferous tubule sections from E13.5 and P0 testis are stained for DDX4, H3K4me3 and DAPI, germ cells are indicated by white arrowheads. See also Figure S2.

Among these activated genes were *Dnmt3l* ^28,29^ (Fig. 2E), *Miwi2* ^30–33^, and *Morc1* ^27,34^ (Fig. S2B), all of which are known to function in gonocytes. The accuracy of our RNA-seq analysis was also supported by the fact that PGC marker genes, as well as pluripotency genes, were downregulated from E13.5 (Fig. S2B), while spermatogonia marker genes were upregulated at P3 (Fig. 2E, S2B), as expected. It also revealed the transiently low expression of *Cdk1* ^35^ (Fig. 2E), illuminating the mitotic cell cycle arrest known as a specific feature of gonocytes ^36^. In addition to the above genes, we unexpectedly observed the upregulation of spermatogenesis-related genes known to function in the adult testis, such as *Meiosin* ^37,38^, *Izumo1* ^39,40^, *Tex101* ^41–43^ (Fig. 2E). Moreover, the transiently activated genes were involved in various cellular pathways and physiological events, including developmental maturation, cognition and the immune system (Fig. 2F).

To better describe the chromatin state during the gonocyte middle stage, we investigated the nuclear signal of H3K4me3, a histone modification correlated with transcriptional activity. The histone modification was enhanced specifically in germ cells at P0, implying that transcription becomes permissive at this stage (Fig. 2G). Moreover, the chromosomes did not exhibit the typical condensed structure observed in meiotic prophase, indicating that meiosis and subsequent spermatogenesis are not induced at this stage (Fig. 2G). These findings suggested that a subset of genes is ectopically expressed in embryonic germ cells, yet this does not lead to aberrant differentiation in such a cellular environment.

Revealing the transient activation of genes in gonocytes, we wondered how it was associated with *de novo* DNA methylation. We sorted genes by their expression levels at P0 and profiled their promoter DNA methylation levels (Fig. S2C). This analysis implied that the elevated expression levels may play a role in shaping the DNA methylation landscape at promoters in gonocytes.

### H3K27me3 Global Erasure Coincides with Gene Upregulation in Gonocytes

Since the molecular mechanism governing the transient gene activation was unlikely to be mediated by *de novo* DNA methylation, it prompted us to investigate the role of other epigenetic marks in this phenomenon. H3K27me3, deposited by Polycomb repressive complex 2 (PRC2), plays a crucial role in gene regulation by mediating transcriptional repression. PRC2-mediated regulation is also indispensable for various mammalian developmental processes, including spermatogenesis ^44–46^. To quantitatively assess the levels of H3K27me3 across the genome, we performed spike-in chromatin immunoprecipitation followed by sequencing (spike-in ChIP-seq) ^47–50^ at different stages in gonocytes (Fig. S3A). Strikingly, H3K27me3 showed a remarkable genome-wide erasure after E13.5, reaching its lowest level at P0 in both genic and intergenic regions, then recovering slightly at P3 (Fig. 3A).

**Fig. 3:**
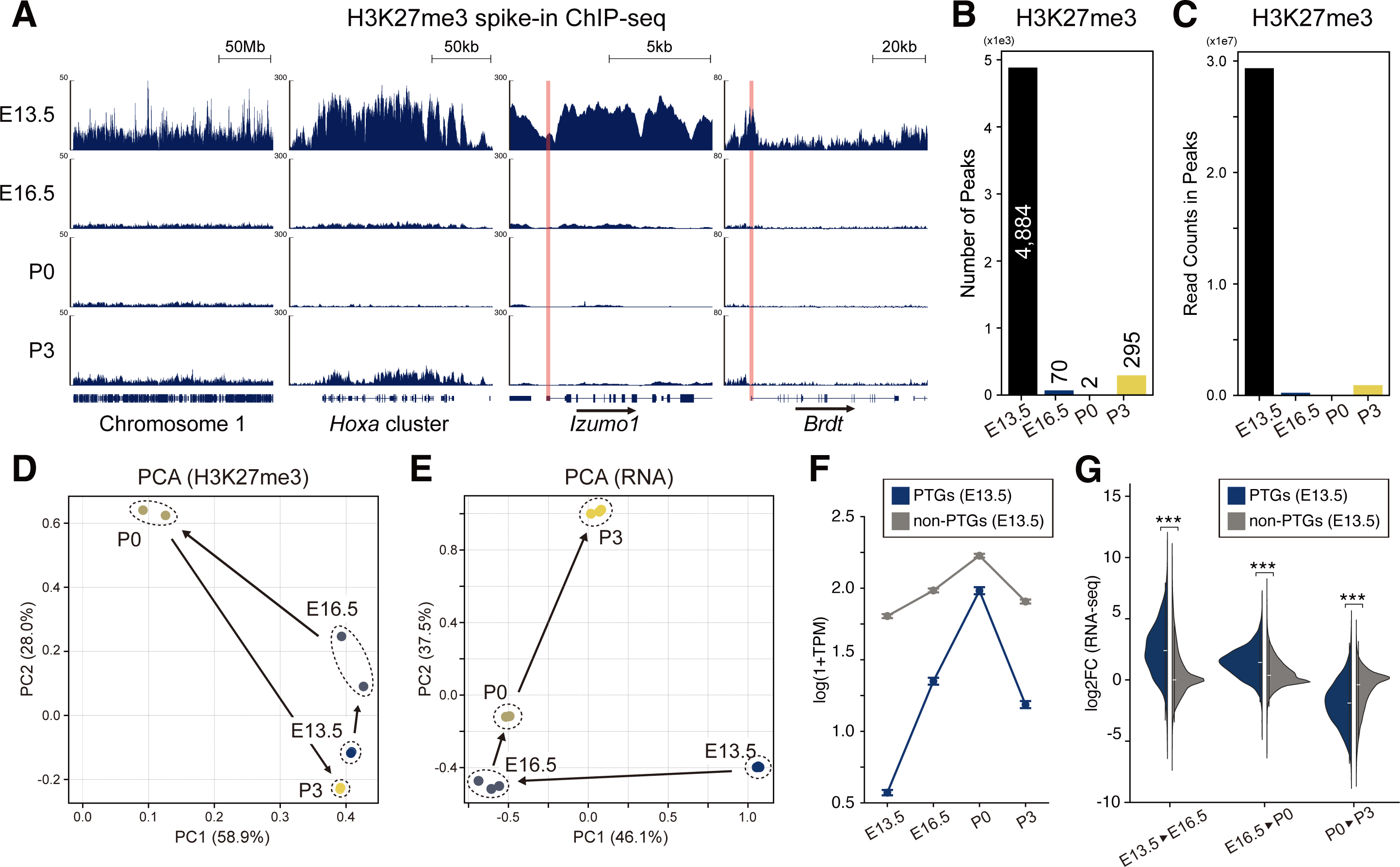
H3K27me3 Global Erasure Coincides with Gene Upregulation in Gonocytes. (**A**) Genome browser representations of H3K27me3 spike-in ChIP-seq data from E13.5 to P3 illustrating Chromosome 1 and gene regions of *Hoxa* cluster, *Izumo1*, and *Brdt*. Transcription start sites are indicated by red lines. (**B**) Number of H3K27me3 peaks for each stage (see Methods). (**C**) Read counts in all identified peaks at each stage. (**D**) Principal component analysis (PCA) of H3K27me3 enrichment at promoter regions (TSS +/-500bp) during gonocyte period (**E**) PCA of normalized expression levels (TPM) of all protein-coding genes during gonocyte period. (**F**) Gene expression levels of E13.5 Polycomb target genes (PTGs, n=2,701) and non-PTGs (n=19,145) at each gonocyte stage. (**G**) Gene expression log2Foldchanges of PTGs and non-PTGs during each period. See also Figure S2, S3.

This erasure was observed not only for conventional PRC2 targets, such as *Hox* genes clustered in four major genomic regions (*Hox a-d*), but also for unconventional targets such as *Izumo1* ^39^ and *Brdt* ^51,52^, known as male germline-specific genes necessary for male fertility functioning in adult testis (Fig. 3A). Notably, the H3K27me3 loss on these two male germline-specific genes was in parallel with their ectopic expression (Fig. 2E, S2D). *Brdt* encodes a transcription factor required for the activation of meiosis-specific genes in adult testis ^53^. BRDT protein was observed in gonocyte nuclei at P0 but not at P3 (Fig. S2E). Moreover, chromatin proteome ^54^ analysis revealed that BRDT transiently bound to gonocyte chromatin, supporting its ectopic expression even at the protein level in embryonic germ cells (Fig. S2F).

Detailed analysis revealed a prominent reduction in both the number and intensity of H3K27me3 peaks from E13.5, followed by a slight recovery at P3 (Fig. 3B, C) (see Methods). Principal Component Analysis (PCA) of the H3K27me3 genome distribution illustrated a U-shaped trajectory, with a gradual change from E13.5 to P0 and a sharp reversal from P0 to P3 (Fig. 3D), suggesting a drastic shift in H3K27me3 levels during the middle stage of gonocyte. The PCA of transcriptomes from E13.5 to P3 also showed a sharp turn at E16.5, followed by gradual changes thereafter. As a result, the P3 transcriptome was reorganized compared to that at E13.5 (Fig. 3E), reflecting the ongoing differentiation of gonocytes into spermatogonia.

To investigate the relationship between transcript upregulation and H3K27me3 removal in each gene, Polycomb target genes (PTGs) were defined based on H3K27me3 density within promoter regions in E13.5 gonocytes (see Methods). Henceforth, we referred to these genes as “PTGs (E13.5)”. PTGs (E13.5) (n=2,701) showed pronounced upregulation from E13.5, followed by marked downregulation from P0 compared to other genes, suggesting a correlation between H3K27me3 elimination and gene activation (Fig. 3F, G). Hereby, we proposed that the global erasure of the repressive histone mark H3K27me3 may underpin the widespread gene upregulation in gonocytes, which has a negative correlation with *de novo* DNA methylation.

### PRCs Elimination from Chromosomes Aligns with Erasure of Repressive Histone Marks

Since we suspected that the global erasure of H3K27me3, accompanied by transient gene activation, may have a role in regulating the receptivity of *de novo* DNA methylation, we investigated the factors responsible for such widespread loss of H3K27me3. At the transcript level, subunits of canonical PRC2, *Ezh2*, *Suz12*, and *Rbbp4/7* ^55–58^, were downregulated during the gonocyte middle stage (Fig. 4A). Some subunits of noncanonical PRC2 were also markedly downregulated (Fig. S4A), while *Ezhip* ^59^, *Eed*, and *Ezh1* were upregulated (Fig. 4A). To reveal the abundance of PRC2 binding to chromosomes, we conducted CUT&RUN ^60–62^ for EZH2, which is an enzymatic core of PRC2. Intriguingly, the number of EZH2 peaks and the signal intensity within these peaks drastically decreased at P0 compared to E13.5 (Fig. 4B, S4B, C), showing the pervasive loss of EZH2 binding from gonocyte chromatin. Accordingly, we supposed that the PRC2 loss of binding may serve as the trigger of H3K27me3 global loss. However, depletion of *Ezh2* from gonocytes at E13.5, which leads to loss of H3K27me3 on the genome ^63^, did not result in upregulation of PTGs as drastic as that observed in the middle stage of gonocyte (Fig. 3G, 4C). This implied that some pathways other than PRC2 may compensate for gene repression in *Ezh2* conditional knockout (cKO) gonocytes at E13.5.

**Fig. 4:**
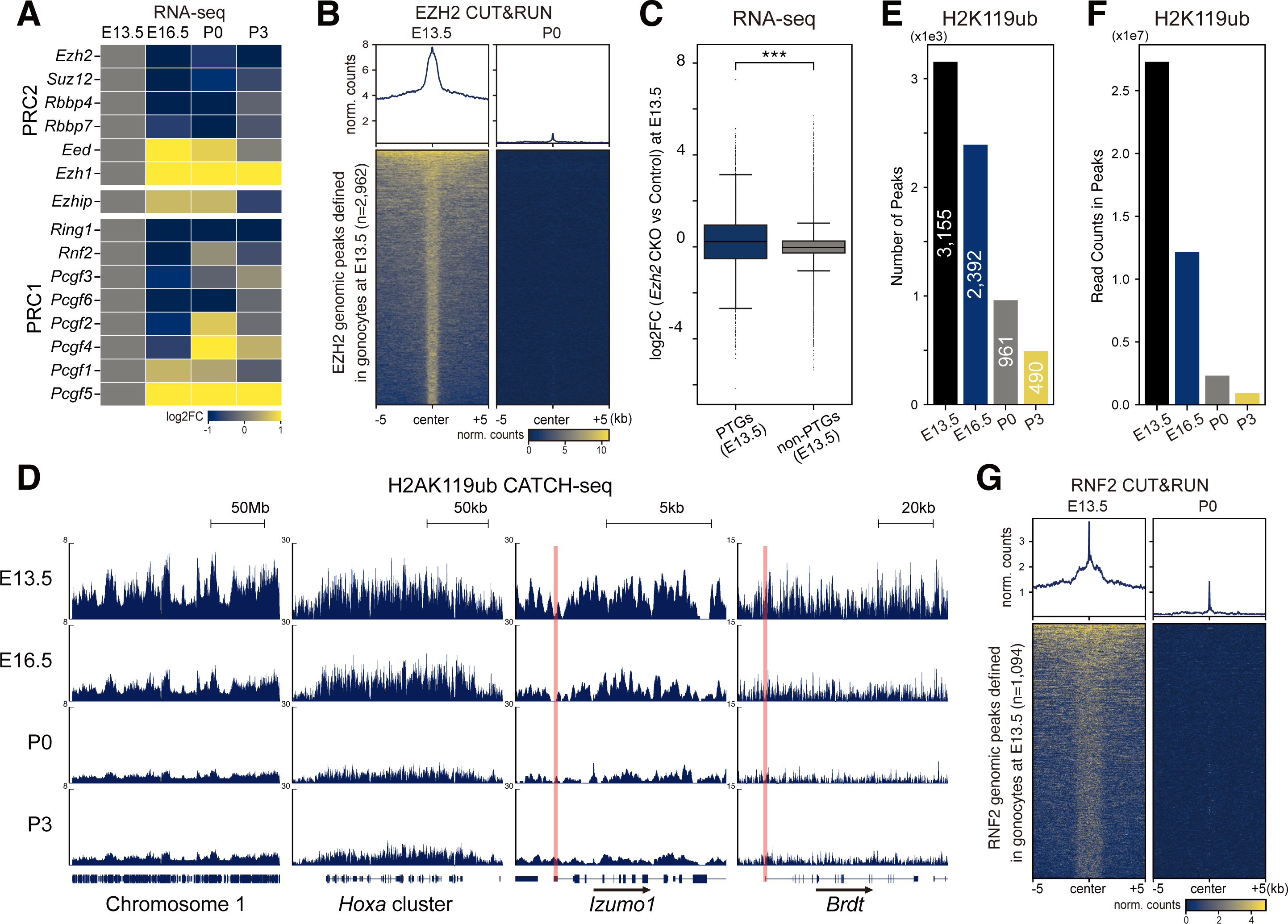
PRCs Elimination from Chromosomes Aligns with Erasure of Repressive Histone Marks. (**A**) Heatmap showing log2FoldChanges between gene expressions of each stage compared with E13.5, core subunit genes of PRC2 and PRC1, as well as *Ezhip* are represented. (**B**) The intensity of EZH2 genomic binding levels at E13.5 and P0 on peaks defined at E13.5. (**C**) Gene expression log2Foldchanges of E13.5 PTGs (n=2,701) and non-PTGs (n=19,145) in E13.5 male PGCs of *Ezh2* cKO and control mouse. (**D**) Genome browser representations of H2AK119ub CATCH-seq data from E13.5 to P3 illustrating Chromosome 1 and gene regions of *Hoxa* cluster, *Izumo1*, and *Brdt*. Transcription start sites are indicated by red lines. (**E**) Number of H2AK119ub peaks for each stage (see Methods). (F) Read counts in all identified peaks at each stage. (G) The intensity of RNF2 genomic binding levels at E13.5 and P0 on peaks defined at E13.5. See also Figure S4.

Since PRC1 has been reported to have an overlapping function with PRC2 in terms of gene repression, we suspected that PRC1 may also be involved in the silencing of the PTGs (E13.5) in gonocytes. Most of the PRC1 core subunits and some noncanonical PRC1 subunits were downregulated from E13.5 (Fig. 4A, S4A). Moreover, quantitative CATCH-seq ^64^ for H2AK119ub at different gonocyte developmental stages (Fig. S3B) revealed that H2AK119ub existed on the genome at E13.5 (Fig. 4D), which may contribute to gene repression at this time point and account for the non-drastic upregulation of PTGs (E13.5) in *Ezh2* mutant gonocytes (Fig. 4C). It also revealed a reduction in H2AK119ub from E13.5, reaching its lowest level at P0 and persisting until P3 (Fig. 4D). While the number and density of H2AK119ub peaks showed a continuous decrease from E13.5 to P3 (Fig. 4E, F). Notably, though the reduction in H2AK119ub was modest and gradual compared to that of H3K27me3, the patterns of their global loss were markedly similar (Fig. 3A, 4D), suggesting their interdependencies of genomic localization as previously reported ^55^.

CUT&RUN for PRC1 core subunit RNF2 revealed a significant reduction in its genome binding level at P0 (Fig. 4G, S4D, E), consistent with the minimal level of H2AK119ub enrichment at that time (Fig. 4D). This epigenetic reorganization, coupled with the H3K27me3 global erasure, putatively contributed to a permissive chromatin state for transcription, which could explain the more pronounced upregulation during the gonocyte middle stage than that in E13.5 *Ezh2* cKO gonocytes (Fig. 3G, 4C). Thus, we concluded that not only PRC2 and H3K27me3, but PRC1 and H2AK119ub underwent erasure across the gonocyte chromatin, representing a novel pattern of epigenetic reprogramming, which, to our knowledge, has not been previously reported in the mouse life cycle.

### Housekeeping Genes and Developmental Genes Avoid *De Novo* DNA Methylation

Thus far, we have observed the loss of PRCs from gonocyte chromatin along with the upregulation of diverse genes. To investigate how PRC2-dependent gene regulation affects *de novo* DNA methylation, we conducted a gene-by-gene analysis using all protein-coding genes and found that gene promoters with a comparably high EZH2 binding level (RPM > 1.5) at E13.5 were resistant to *de novo* DNA methylation (Fig. 5A, lower right). GO analysis using EZH2-enriched genes revealed an enrichment of various developmental processes and signaling pathway-associated terms (Fig. S5A). On the other hand, unmethylated genes with low EZH2 binding level at their promoters (RPM ≤ 1.5) (Fig. 5A, lower left) were mostly housekeeping genes involved in cell division, histone modification, ncRNA processing, and DNA repair (Fig. S5B). Hence, we hereafter termed these two groups of genes *Developmental Genes* and *Housekeeping Genes*, respectively (Fig. S5C). We also extracted the hypermethylated (>80%) genes (*High DNAme Genes*) as a control for the subsequent analysis (Fig. S5C).

**Fig. 5:**
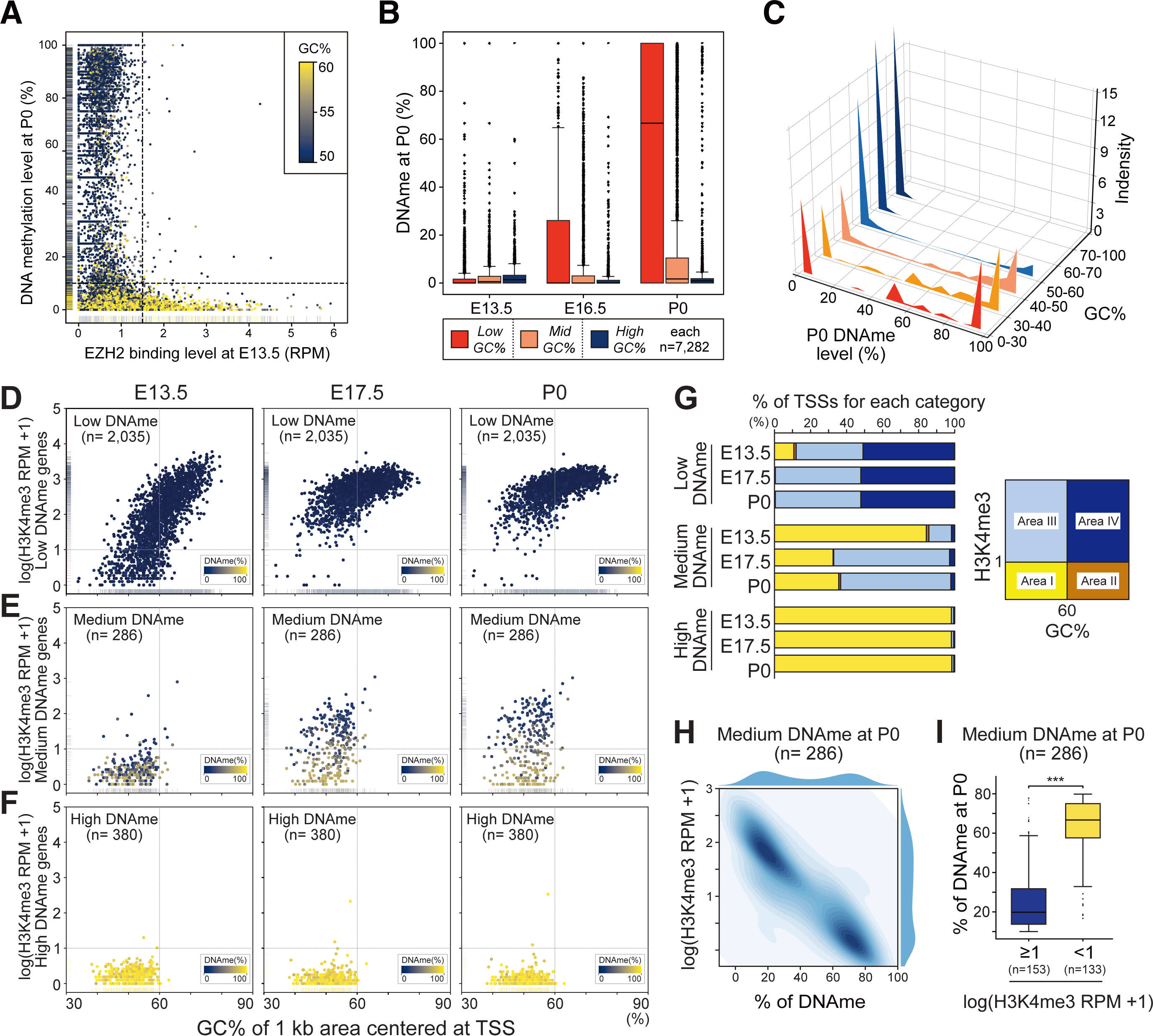
Combination of GC% and H3K4me3 Density Robustly Predicts *De Novo* DNA Methylation Levels at Gene Promoters. (**A**) Scatter plot describing the correlation between E13.5 EZH2 genomic binding level and P0 DNA methylation in promoters. Scatters are annotated with a colormap based on promoter GC%. (**B**) Boxplots showing P0 promoter DNA methylation levels of three gene groups defined in Fig. S5F. (**C**) Density plot showing analysis of P0 DNA methylation levels in six groups categorized by GC%. (**D**) Scatter plot illustrating the correlation between GC% and H3K4me3 density level in promoters of Low DNAme genes, as defined by P0 DNAme level ranging from 0% to 10%. Each dot of gene is annotated with a colormap based on P0 DNAme level. (**E**) As in (**D**), the gene sets with Medium DNAme genes as defined by P0 DNAme level ranging from 10% to 80% are plotted. (**F**) As in (**D**), the gene sets with High DNAme genes as defined by P0 DNAme level ranging from 80% to 100% are plotted. (**G**) The two-dimension area of scattered plots shown in (**D**), (**E**), and (**F**) are divided into four sections by thresholds 60% of GC% and one of log(H3K4me3 RPM + 1) as indicated by the illustration figure on the right. The proportion of the gene numbers in each area is shown in the bar graph. (**H**) Contour plot and density plot illustrating the correlation between P0 DNA methylation and P0 H3K4me3 density level on genes in the P0 Medium DNAme group. (**I**) Boxplots showing P0 DNA methylation levels for two gene sets within the P0 Medium DNAme group, categorized by their H3K4me3 density being higher or lower than a threshold of one for log(H3K4me3 RPM + 1). See also Figure S5.

*High DNAme Genes* progressively acquired *de novo* DNA methylation during the gonocyte period, while *Developmental Genes* and *Housekeeping Genes* remained unmethylated (Fig. S5D left). Interestingly, *Developmental* and *Housekeeping Genes* showed higher expression levels than *High DNAme Genes* already from the early stage of gonocytes (Fig. S5D right). Since both gene groups, *Developmental* and *Housekeeping Genes*, achieved pronounced high levels of gene activity during the gonocyte middle stage, we suspected that such high gene activities in gonocytes may help them avoid *de novo* DNA methylation.

### GC% is Not the Sole Determinant of DNA Methylation Levels

Since some sequence features correlate with DNA methylation ^65^, we searched for nucleotide characteristics of promoters that would determine their *de novo* DNA methylation level. Overall, the promoters with low GC% were more likely to be methylated compared with those with higher GC% (Fig. 5B, S5E, F). Moreover, *Developmental Genes* and *Housekeeping Genes*, defined above (Fig. S5C), showed higher GC%. These investigations were in parallel with previous findings indicating the negative correlation between GC% and steady-state DNA methylation level ^66,67^.

However, detailed analysis revealed certain promoters with low GC% avoided *de novo* DNA methylation (P0 DNAme level 0-10%), while others acquired high DNA methylation level (90-100%) (Fig. 5C). This bimodal distribution pattern of the *de novo* DNA methylation level at these low GC% promoters suggested that GC% alone is not sufficient to predict the extent to which the promoters acquire *de novo* DNA methylation.

### Combination of GC% and H3K4me3 Density Robustly Predicts *De Novo* DNA Methylation Levels at Gene Promoters

Since the level of H3K4me3 generally correlates with transcriptional activity ^68,69^ and was enhanced during the middle stage of gonocyte (Fig. 2G), we next investigated how the dynamic changes in H3K4me3 density may correlate with the level of *de novo* DNA methylation on promoters in gonocytes. Specifically, PTGs (E13.5) (defined in Fig. 3F) were categorized into three groups by their DNA methylation level at P0, then the density of H3K4me3 was profiled alongside GC% in their promoters from E13.5 to P0 (Fig. 5D-F). For all three groups, H3K4me3 experienced an increase from E13.5 to P0, and its density showed a negative correlation with DNA methylation level, as reported in earlier studies ^70–74^. Importantly, H3K4me3 density at promoters with relatively low GC% (< 60%) experienced a more remarkable increase (Fig. 5D-F), indicating that the activation of these genes was more pronounced. The increases in H3K4me3 also differed significantly among the three groups. Compared with the High DNAme group (Fig. 5F), which showed little increase in H3K4me3 at the promoters, the Medium and Low DNAme groups exhibited a mild and more pronounced increase in promoter H3K4me3, respectively (Fig. 5D, E). This illustrated that the increase in H3K4me3 density is highly associated with the establishment of *de novo* DNA methylation in gonocytes (Fig. 5D-F).

Notably, the H3K4me3 densities of the Medium DNAme group were comparable across genes at E13.5 but showed significant variation in the degree of elevation by P0 (Fig. 5E). Assuming H3K4me3 density as log(RPM + 1) = 1 and GC% as 60% as putative thresholds that might specify the likelihood of *de novo* DNA methylation, all subplots were divided into four areas to analyze the correlation between quantitative changes in H3K4me3 density, intrinsic GC%, and *de novo* DNA methylation (Fig. 5G). Almost all promoters in the Low DNAme group (2,026 out of 2,035) showed an increase in H3K4me3 density during the gonocyte period, surpassing the putative threshold observed at P0. In contrast, no significant changes in H3K4me3 density were found in the High DNAme group with 378 of 380 promoters remaining below the threshold for H3K4me3 density (Fig. 5G). The Medium DNAme group represented a transient state, showing two populations with different DNA methylation levels around the H3K4me3 boundary (Fig. 5D-F). The density plots of H3K4me3 for genes within the Medium DNAme group displayed a bimodal distribution, representing distinct susceptibility to *de novo* DNA methylation: gene with increased H3K4me3 density corresponded to lower levels of *de novo* DNA methylation, and those with minimal H3K4me3 increase were linked to higher *de novo* DNA methylation levels (Fig. 5I).

These findings suggested that epigenetic H3K4me3 levels and genetic GC% serve as important quantitative factors determining whether a gene acquires or resists *de novo* DNA methylation. Then, we proposed a model by which most developmental genes and housekeeping genes avoid *de novo* DNA methylation in gonocytes (Fig. S5G). In this model, the global reduction of H3K27me3 and H2AK119ub, along with PRC1 and PRC2 loss of binding, putatively create a permissive transcriptional state during the gonocyte middle stage, potentially leading to higher gene activity and H3K4me3 density, which could contribute to shaping the distinct DNA methylation landscapes across gene promoters (Fig. S5G).

### SSC-specific Chromatin State Is Established through H3K27me3 Restoration from Gonocyte Onward

Having profiled dramatic epigenetic reorganization during the gonocyte period, we wondered how these events may affect the chromatin state in the subsequent developmental stage. Interestingly, the enrichment of H3K27me3 at promoters was restored in undifferentiated spermatogonia at P7 (Undiff Spg (P7)) after its elimination at P0 (Fig. 6A). Despite the similar distribution of H3K27me3 peaks between P7 undifferentiated spermatogonia and E13.5 gonocytes across the genome (Fig. S6A, B), the compositions of Polycomb target genes (PTGs) at these two stages were clearly different (Fig. 6B). For example, the E13.5 PTGs included genes crucial for gonocyte development, such as *Dnmt3b* and *Dnmt3l* ^28,29^, while Undiff Spg (P7) PTGs comprised genes involved in spermatogonia maintenance and subsequent spermatogenesis, such as *Dmrt1* ^75^ and *Kit* (c*-Kit*) ^76,77^ (Fig. 6C). Moreover, at several conventional Polycomb target sites, including *Wnt* clusters, *Hox* clusters, and *Sox* clusters (Fig, 6C), the density of H3K27me3 was restored in spermatogonia, indicating the presence of epigenetic memory guiding the restoration of H3K27me3 at these sites. Since H2AK119ub was partially retained on the genome when H3K27me3 was almost erased in gonocytes and it was previously reported to precede the deposition of H3K27me3 ^64,78^, we supposed H2AK119ub may serve as the epigenetic memory. Importantly, H3K27me3 was largely absent in undifferentiated spermatogonia of PRC1 cKO adult mouse ^12^ (Fig. 6D), suggesting that PRC1-mediated H2AK119ub contributes to the redeposition of H3K27me3 on the genome. Of note, PRC1 cKO mice are unable to produce functional sperm and ultimately cause infertility ^11^, indicating that the restoration of H2AK119ub and H3K27me3 following the gonocyte period is indispensable for subsequent spermatogenesis.

**Fig. 6:**
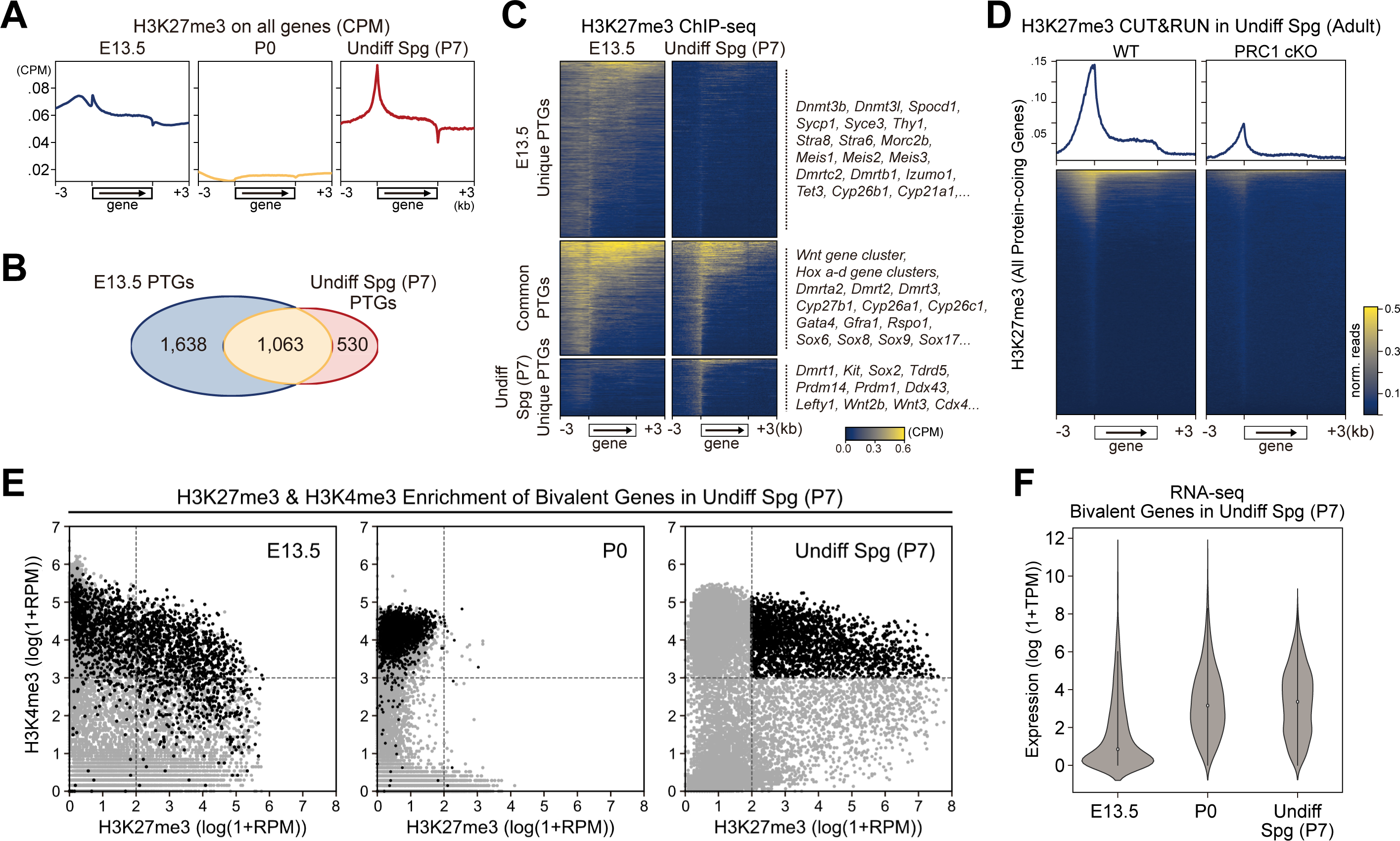
SSC-specific Chromatin State Is Established through H3K27me3 Restoration from Gonocyte Onward. (**A**) Lineplots of the average H3K27me3 intensity on all protein-coding genes in E13.5 and P0 gonocytes as well as P7 undifferentiated spermatogonia. (**B**) Venn diagram of Polycomb target genes (PTGs) defined in E13.5 gonocytes and P7 undifferentiated spermatogonia. (**C**) Heatmap of H3K27me3 intensity on PTGs defined in E13.5 gonocytes and P7 undifferentiated spermatogonia. PTGs uniquely identified in either stage, as well as those commonly identified in both stages are listed. (**D**) Heatmap of H3K27me3 intensity on all protein-coding genes in WT and PRC1 cKO adult undifferentiated spermatogonia. (**E**) Scatterplot illustrating H3K27me3 and H3K4me3 enrichment within promoters in E13.5 and P0 gonocytes, as well as P7 undifferentiated spermatogonia. Black scatter points indicate bivalent genes identified in P7 undifferentiated spermatogonia. (**F**) Expression levels of bivalent genes in P7 undifferentiated spermatogonia at each stage. See also Figure S6.

Bivalent chromatin, characterized by the simultaneous presence of H3K4me3 and H3K27me3, plays a crucial role in maintaining a set of developmental genes poised in SSCs ^16,79^. To investigate how bivalent chromatin was constructed during the transition from gonocytes to spermatogonia, we defined bivalent genes in undifferentiated spermatogonia (P7) ^12^ and traced back their epigenetic states (Fig. 6E). This revealed that 40.0% of Undiff Spg (P7) bivalent genes were already pre-set in the bivalent state at E13.5 (Fig. 6E). This bivalent state was lost at P0 but subsequently re-established by P7 (Fig. 6E), identifying the developmental windows for both epigenetic reprogramming and programming. Notably, these Undiff Spg (P7) bivalent genes experienced an increase in H3K4me3 at P0, preceding H3K27me3 restoration (Fig. 6E), and a significant upregulation in their expression levels (Fig. 6F). Thus, we proposed a two-step chromatin priming model for establishing SSC-specific chromatin state at promoters. The first step involves avoiding *de novo* DNA methylation through the dynamic increase in H3K4me3, while the second step entails establishing the bivalent state through the H3K27me3 restoration at hypo-methylated promoters. These findings highlighted a critical time window for the establishment of SSC-specific chromatin state.

### Epigenetic Priming in Gonocytes May Have Long-term Effects on Subsequent Development

The genes activated in gonocytes are highly diverse, including Polycomb targets. Previous analysis implied that germ cell populations display distinct differentiation patterns from the gonocyte middle stage onward ^80^. Therefore, we conducted single-cell RNA-seq analysis (scRNA-seq) using gonocytes to investigate whether the ectopic expression of genes originated from specific cell populations (Fig. 7A). The expression pattern of several marker genes, such as *Cdk1* ^35^, *Dnmt3l* ^28,29^, *Nanog* ^81,82^, and *Gfra1* ^83^, indicated that our scRNA-seq dataset precisely reflected the known developmental processes occurring in gonocytes (Fig. 7A). Several genes associated with adult testis function, such as *Tex101* ^41–43^*, Ropn1l* ^84,85^*, Dpep3* ^86,87^, and *Nme8* ^88–90^, were transiently expressed during the middle stage of gonocyte, exhibiting little intercellular variation (Fig. 7A, S7A). Genes typically expressed in other tissues rather than testis, such as *Cartpt* ^91^, *Serping1* ^92–94^, *Scn8a* ^95,96^, and *Fcer1g* ^97,98^, also showed ectopic expression in gonocytes (Fig. 7A, S7A). Notably, single-cell transcriptomes at each developmental stage showed marked homogeneity (Fig. 7A), revealing that the ectopic expression was not a result of premature or abnormal cell development. These findings aligned with the above suggestion that the genes are pre-activated in gonocytes shaping the epigenetic landscape, resembling the “epigenetic priming” often seen in other cell lineages ^99,100^.

**Fig. 7:**
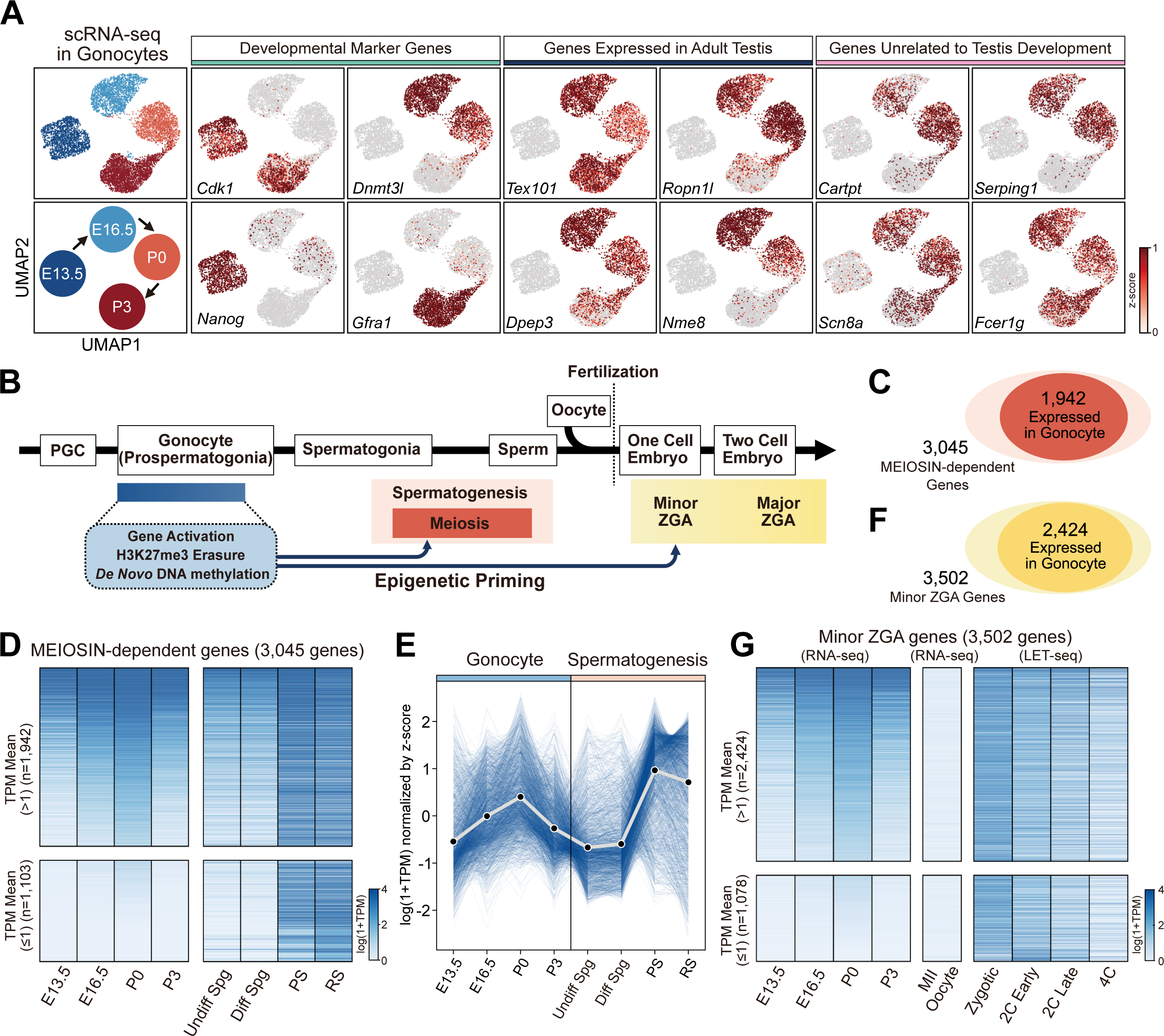
Epigenetic Priming in Gonocytes May Have Long-term Effects on Subsequent Development. (**A**) Gene expression level at single cell level revealed by scRNA-seq data of gonocytes at four stages as described in the leftmost column. Expression patterns of three groups of genes are demonstrated in three panels, respectively. (**B**) Schematic illustrating developmental timeline from PGC to preimplantation embryo, highlighting key developmental processes at each stage. (**C**) Schematic illustrating the numbers of MEIOSIN-dependent genes and the subset expressed in gonocytes. (**D**) Heatmap of MEIOSIN-dependent genes (n=3,045) including both subsets with TPM mean higher (n=1,942) or lower than one (n=1,103) in gonocytes, illustrating their expression changes throughout the gonocyte and subsequent spermatogenesis stages. Undiff Spg: P7 undifferentiated spermatogonia, Diff Spg: P7 differentiating spermatogonia, PS: pachytene spermatocyte, RS: round spermatid. (**E**) Line plot illustrating expression changes in the subset of MEIOSIN-dependent genes that are expressed in gonocytes (n=1,942), illustrating their expression levels throughout the gonocyte and subsequent spermatogenesis stages. (**F**) Schematic illustrating the numbers of Minor ZGA genes and the subset expressed in gonocytes. (**G**) Heatmap of Minor ZGA genes (n=3,502) including both subsets with TPM mean higher (n=2,424) or lower than one (n=1,078) in gonocytes, illustrating their expression levels in gonocytes, MII oocytes, and preimplantation embryos. See also Figure S7.

This prompted us to evaluate the extent to which these ectopically expressed genes are utilized in subsequent developmental stages, for example, spermatogenesis and early embryogenesis in the next generation (Fig. 7B). The progeny cell type of gonocytes is the spermatocyte, which has the potential to initiate meiosis. We focused on a specific group of genes whose expression is downregulated in the adult testes of *Meiosin* knockout mice (8 weeks old) ^37^. Since MEIOSIN is a pivotal transcription factor required for the initiation of meiosis, these MEIOSIN-dependent genes are typically activated during meiosis or later stages of spermatogenesis ^37^. Strikingly, nearly two-thirds of the MEIOSIN-dependent genes (1,942 out of 3,045) were expressed in gonocytes (Fig. 7C, S7B), levels of which were lower than those observed in the adult testis (Fig. 7D, E).

We then focused on genes expressed during the developmental stage immediately following fertilization, known as zygotic genome activation (ZGA), which governs the maternal-to-zygotic transition of the transcriptome and plays a crucial role in preimplantation development ^101–107^. Minor ZGA, preceding major ZGA which begins at the two-cell stage, starts in the zygote when parental DNA methylation is almost maintained ^3,108–110^. We utilized 3,502 Minor ZGA Genes ^107^ and revealed that nearly 70% of them (2,424 out of 3,502) were transiently activated in gonocytes (Fig. 7F, G, and S7B). This suggested that the priming of certain genes in gonocytes may have a broader effect on development that spans to the preimplantation embryo, though transcripts produced from the maternal genome also contribute to these newly synthesized RNAs. Based on these findings, we proposed that epigenetic memory at gene promoters mediated by gene activation in embryonic germ cells may have long-term effects not only on spermatogenesis but also on early embryonic development.

## Discussion

### Shaping DNA methylation Landscapes at Gene Promoters in Gonocytes

In this work, we investigated how *de novo* DNA methylation landscape is established at gene promoters in mouse gonocytes. During this developmental stage, diverse genes accounting for 70% of protein-coding genes, including housekeeping genes and developmental genes, were kept hypomethylated. Unexpectedly, these genes were transiently activated and transcribed during the middle stage of gonocyte. This upregulation was concomitant with the global erasure of repressive histone marks of both H3K27me3 and H2AK119ub, as well as the elimination of the genomic localization of the subunits of PRC1 and PRC2. Such genes showed the accumulation of H3K4me3 over their promoter regions, accompanied by selective resistance to *de novo* DNA methylation. This is consistent with the earlier biochemical evidence showing that DNMT3A and DNMT3L, the methyltransferase and its cofactor involved in *de novo* DNA methylation, only bind to chromatin lacking H3K4 methylation ^70,111^. Furthermore, we revealed that a specific level of H3K4me3 density would work as a robust probe distinguishing whether promoters acquired *de novo* DNA methylation or not. Therefore, we speculated that boosting the basal density of H3K4me3 on a vast majority of protein-coding genes serves as a special strategy to counteract DNA methylation-mediated silencing.

### Unconventional Epigenetic Plasticity in Male Germ Cells

H3K27me3 underwent a global reduction during the gonocyte stage. Although such erasure of H3K27me3 has been rarely seen in the mouse life cycle, a somewhat similar redeposition of H3K27me3 occurs in preimplantation embryos ^112,113^. In the fertilized egg of mouse, H3K27me3 landscapes on parental chromosomes experience reorganization and show a non-canonical genomic pattern persisting until E6.5. Although H3K27me3 on the promoter of many genes is lost through this redistribution, the expression of these genes shows no obvious activation, representing a clear contrast with our observation in gonocytes. The rewiring of H3K27me3 distribution in early embryos was suggested to help reset many gene promoters to a reprogrammed state, though molecular validation has not been performed yet.

During the transition from gonocyte to spermatogonia, the density of H3K27me3 was not only restored but also newly deposited onto stage-specific developmental gene promoters. Notably, the re-establishment of H3K27me3 was dependent on the existence of PRC1, which is consistent with what has been reported in other systems that H2AK119ub acts as a preceding mediator of H3K27me3 ^64,78^. Interestingly, H2AK119ub also showed a global decrease in gonocytes, implying an intricate relationship between the two histone modifications. As the reduction in H2AK119ub was milder and delayed compared to that of H3K27me3, we supposed that the remaining H2AK119ub may serve as an epigenetic memory, guiding the redeposition of H3K27me3 onto the genome.

### Establishing Epigenetic Priming State in Male Germ Cells

By illustrating how gene promoters adopted a bivalent state, which has essential roles in SSC function, we proposed that the epigenetic dynamics in gonocytes contributes to establishing epigenetic priming in SSC. Notably, this epigenetic priming differs from the conventional pattern in at least two aspects. First, in typical cases, primed genes are poised with transcription elongation blocked. In contrast, primed genes in gonocytes are transcriptionally activated. Second, the H3K27me3 redistribution occurring in gonocytes was genome-wide, rather than being limited to promoter regions as seen in conventional epigenetic priming. These differences may be attributed to the fact that the establishment of epigenetic priming occurs simultaneously with the introduction of DNA methylation in gonocytes. In other words, maintaining the DNA hypomethylation state at developmental gene promoters in gonocytes would be a prerequisite step for the establishment of epigenetic priming in male germ cells, especially spermatogonial stem cells ^20,79^. It will be the subject of future research to reveal the key transcription factor(s) responsible for this epigenetic priming.

### Functional Interaction between PRC2 and Cell Cycle Arrest

Male germ cells experience a mitotic cell cycle arrest at G_0_/G_1_ in gonocytes and re-enter the cell cycle afterward to further differentiate ^114^. Of note, the initiation of genome-wide erasure of H3K27me3 in gonocytes coincides with the onset of the cell cycle arrest. This correlation between cell cycle and Polycomb repressive complex has also been reported in cancer cells, where downregulation of PRC2 subunits leads to H3K27me3 decrease and proliferation inhibition, characterized by the increase in the number of cells arrested at G_0_/G_1_ phase ^115–117^. On the other way around, cancer cells blocked at G_1_ phase express less EZH2 protein ^118^. Accordingly, cell cycle arrest and the downregulation of PRC2 subunits may be correlated interdependently in gonocytes. Since PRC2 plays an indispensable role in germ cell development, it is challenging to verify the effect of PRC2 depletion on the cell cycle using a normal conditional knockout system, due to the limitation of time resolution. Recently, the auxin-inducible degron (AID) system has enabled the rapid and reversible depletion of PRC2 subunit proteins in cultured cells ^119^, and its application *in vivo* might facilitate future exploration of this important subject.

In summary, we profile the temporal dynamics of the epigenome in gonocytes and reveal how genes are regulated through epigenetic reprogramming. We previously showed that gonocyte chromatin adopts a relaxed structure, as evidenced by the transient loss of higher-order chromatin conformations and the reactivation of myriad transposons ^21^. The current study shows that the global elimination of repressive histone marks correlates with a transient upregulation of thousands of genes. These phenomena indicate that the entire chromatin undergoes reprogramming so that the genetic circuit becomes rewired in gonocytes. This dynamics may contribute to preventing epimutation and aberrant epigenetic information from being passed on to the next generation with potentially detrimental effects. Future studies should reveal the key factors triggering such genome-wide events and whether similar events also occur during mammalian oogenesis.

## Methods&Materials

### Animal Care and Use

All animal procedures were approved by the Institutional Safety Committee on Recombinant DNA Experiments and the Animal Research Committee of The University of Tokyo. All animal experiments were performed under the guidelines for animal experiments at The University of Tokyo.

### Isolation of Germ Cells from Testes

Testes were obtained from E13.5, E16.5 Mvh-Venus TG embryos and P0, P3 newborn Mvh-Venus TG pups ^120^. After removing the tunica, dissociation buffer [500μL Dulbecco’s modified eagle medium (DMEM), 10μL fetal bovine serum (FBS), 7.5μL of 100 mg/mL hyaluronidase (Tokyo Kasei, Japan, H0164), 2.5μL of 10 mg/mL DNAse (Sigma, D5025-150kU), 10μL of 100 mg/mL collagenase (Worthington, CLS1), and 25μL of 14,000 U/mL recombinant collagenase (Wako, 036-23141)] was applied at 37°C for 20 min. Then testicular cells were completely dissociated by rigorous pipetting and resuspended in 2% FBS/PBS. Subsequently, Venus-positive cells were isolated by fluorescence-activated cell sorting using a FACS Aria III (BD) after adding propidium iodine to select viable cells.

### Histology and Immunostaining

Testis were fixed with 4% paraformaldehyde (PFA) for one hour at 4°C with gentle inverting, then washed in PBS and incubated stepwise in 10 %, 20% and 30% sucrose in PBS at 4°C. The samples were then mounted in OCT mounting medium (VWR) and stored at -80°C. The mounted samples were sectioned for 10 µm using cryostat (Epredia HM525 NX). The cryosections were autoclaved in 10mM citric acid buffer at 105°C for 15min and washed with PBS-T three times. Sections were blocked with 3% skim milk in PBS-T at RT for 30min and incubated with primary antibodies in 3% skim milk in PBS-T at RT for 60min. Then, sections were washed with PBS-T at RT for 10min three times and incubated with secondary antibodies in 3% skim milk in PBS-T at RT for 60min. Mouse anti-DDX4 (1:1000, ab27591), Mouse anti-BRDT (1:50, sc-515674), Rabbit anti-H3K4me3 (1:1000, ab8580) primary antibodies, as well as Alexa Fluor 555 Goat anti-Rabbit IgG (H+L) (1:1000, a21428) and Alexa Fluor 488 Goat anti-Mouse IgG1 (1:1000, a21121) secondary antibodies were used in this study. Finally, the sections were washed with PBS-T at RT for 10min three times and mounted with VECTASHIELD Mounting Medium with DAPI (Vector Laboratories). Images were obtained using LSM980 laser scanning confocal microscope (Carl Zeiss).

### scRNA-seq Library Construction

5,000 germ cells isolated respectively from E13.5, E16.5, P0 and P3 testis were used for experiment. Library construction was performed using the Chromium GEM Single Cell 3’ Reagents kit v3.1 (10x Genomics) in accordance with the manufacturer’s protocol. Sequencing was performed on a Novaseq 6000 (Illumina) and the read length configuration was 28×8×91 cycles for Read 1, Index, and Read 2, respectively. Cell Ranger Single Cell Software Suite 3.1.0 was used to perform sample de-multiplexing, barcode processing, and single cell gene counting.

### Spike-in ChIP-Seq Library Construction

Venus-positive testicular germ cells (2 × 10^4^) were fixed with 1% formaldehyde for 10min. Cells were resuspended in Swelling buffer [20mM Hepes (pH 7.9), 1.5mM MgCl_2_, 10mM KCl, 0.1% NP-40, and 1mM DTT] and incubated on ice for 20min, followed by centrifugation to remove supernatant. Pelleted nuclei were resuspended in 1×shearing buffer (Covaris, 520154) and fragmented using a sonicator (BRANSON, SFX150). Fragmented products diluted with RIPA buffer [50mM Tris-HCl (pH 8.0), 150mM NaCl, 2mM EDTA (pH 8.0), 1% NP-40, 0.5% sodium deoxycholate, and 0.1% SDS] were pre-cleared with Dynabeads-ProteinG (Thermo Fisher, 11201D) for 1h at 4°C. 2μL of 1000-fold SNAP-ChIP solution (EpiCypher, SNAP-ChIP® K-MetStat Panel) was added to the pre-cleared solution and whole cell extract (WCE) was collected. Immunoprecipitation was performed with 2μL of 1μg/μL anti-H3K27me3 antibody on Dynabeads M-280 Sheep anti-mouse IgG (Thermo Fisher, 11201D) overnight at 4 °C. Beads were washed with low buffer [0.1% SDS, 1% Triton X-100, 2mM EDTA (pH 8.0), 150mM NaCl, and 20mM Tris-HCl (pH 8.0)] four times and then with high buffer [0.1% SDS, 1% Triton X-100, 2mM EDTA (pH 8.0), 500mM NaCl, and 20mM Tris-HCl (pH 8.0)] once. Both IP and WCE samples were incubated with the suspension of direct elution buffer [10mM Tris-HCl (pH 8.0), 5mM EDTA (pH 8.0), 300mM NaCl, and 0.5% SDS] at 65°C for 15min to elute products from beads. Samples were then treated with proteinase K for 6h at 37°C, followed by reversal of cross-linking overnight at 65°C. DNA was extracted by EtOH precipitation and the library was constructed with a QIAseq Ultralow Input Library Kit (Qiagen, 180492) following the manufacturer’s instructions.

### CATCH-Seq Library Construction

CATCH-seq is an improved method of the ultra-low-input native ChIP-seq (ULI-NChIP) ^121^, and the original protocol of CATCH-seq was described in the previous report ^122^. We have made some changes in CATCH-seq to improve usability ^64^. Pellets from 10,000 germ cells were lysed to 9 µL of Nuclei EZ lysis buffer (Sigma, NUC-101) supplemented with a complete EDTA-free protease inhibitor cocktail and 1 mM phenylmethanesulfonyl fluoride. The samples were added with 1 µL of the lysis buffer containing 2,000 Drosophila melanogaster S2 cells (Thermo Fisher Scientific, R69007) for spike-in normalization purposes. Then, 1 µL of a 1% Triton X-100 (Merck 93443) and 1% deoxycholate (Nacalai, 10712-54) mixture solution was added to the samples, which then sat on ice for 5 min. The chromatin was fragmented by 2 U/µl MNase (M0247S, NEB) in 1xMNase buffer supplemented with 1% PEG6000 (Hampton Research, HR2-533) and 2 mM DTT (Nacalai) at 37°C for 7.5 min. The MNase reaction was stopped by adding 1/10 volume of 100 mM EDTA and 1/12 volume of the 1% Triton X-100 and 1% deoxycholate mixture, then the samples were rested on ice for 15 min. The chromatin lysates were then added with freshly prepared immunoprecipitation buffer ^121^, and 5% volume was kept for input library construction. To the rest of the lysate, 30 ng of annealed I-SceI carrier DNA was added ^122^. The forward and reverse strands of the carrier DNA were as follows: /5AmMC6/Gtagggataacagggtaattagggataacagggtaattagggataacagggtaattagggataacagggtaattagggataac agggtaattagggataacagggtaat*c/3AmMO/ and /5AmMC6/Gattaccctgttatccctaattaccctgttatccctaattaccctgttatccctaattaccctgttatccctaattaccctgttatccctaat taccctgttatcccta*c/3AmMO/, respectively, where asterisks represent phosphorothioate bonds. The oligos were synthesized by Integrated DNA Technologies. For each immunoprecipitation reaction, 0.5 µL of rabbit anti-H2AK119ub1 (Cell Signaling Technologies, 8240) conjugated to precleared Dynabeads Protein A (Thermo Fisher Scientific, 10006D) and G (Thermo Fisher Scientific, 10007D) mixture was used. The specificity of the antibody was validated in previous studies ^78^. After immunoprecipitation at 4°C overnight, the chromatin-Dynabeads were washed twice each with the low and high salt wash buffers, and the chromatin was eluted in the freshly prepared ChIP elution buffer at 65°C for 1 hour ^121^. DNA was recovered by phenol-chloroform extraction followed by ethanol precipitation. Adaptor ligation was performed by NEBNext Ultra II DNA Library Prep Kit for Illumina (E7645, NEB) in a half scale of the manufacturer’s instruction, and the libraries were purified by 1.8x SPRIselect beads (B23318, Beckman Coulter). The DNA was amplified by KAPA Hifi 2X mater PCR mix (KK2605) for 13-15 PCR cycles with dual indexing primers (NEBNext Multiplex Oligos for Illumina, E6440). After purification with 0.9x SPRIselect beads, the samples were digested by I-SceI (5 U/µl, NEB, R0694) at 37°C for 2 hrs followed by heat inactivation at 65°C for 20 min and purified by 0.9x SPRIselect beads. The second amplification was not performed. The libraries were sequenced on HiSeq X Ten platform.

### CUT&RUN Library Construction

20,000 cells for each replicate were immobilized to Concavalin-A (ConA) beads and incubated with 1.5μL EZH2 antibody (rabbit, CST #5246) or 1.5μL RNF2 (RING1B) antibody (rabbit, mAb #5694) overnight at 4°C. Beads were washed with Digitonin Buffer three times and incubated with 1.5μL second antibody (anti-rabbit guinea pig, antibodies-online ABIN101961) for 1h at 4°C, followed by washing with Digitonin Buffer three times and pAG-MNase for 1h minutes at 4°C. Beads were then incubated with 2μL 100mM CaCl_2_ for 30min on ice after three times of washing and diluted with 100μL 2xSTOP buffer. The supernatant was collected and 2μL 10% SDS together with 2.5μL ProK were added followed by incubation for 1h at 50°C. Then, the supernatant was collected by mixed solutions with 20μL 3M NaOAc and 200μL Phenol/Chloroform/Isoamyl alcohol (25:24:1). 500μL EtOH and 2ul pallet paint were added to the products followed by incubation for 30min at -20°C and pellets were collected after centrifugation. Final products were suspended with 30ul QIAGEN EB buffer (10mM Tris-HCl) and DNA was extracted with NEBNext® Ultra™ II DNA Library Prep Kit for Illumina® (E7645) following the instruction of the manufacturer.

### Proteomics and Data Analysis

Chromatin enrichment for Proteomics (ChEP) was performed as described previously ^54,123^. Briefly, germ cells were fixed with formaldehyde, lysed and digested with RNase. Nuclei were resuspended with 4% SDS buffer and mixed with 8M Urea buffer. Nuclei were washed with SDS buffer and storage buffer followed by ultrasonication for DNA shearing. The samples were boiled to reverse formaldehyde cross-links, reduced, alkylated and digested by phase transfer surfactant-aided digestion method ^124^. Peptides were desalted on a C18-SCX StageTips ^125^ and applied LC-MS/MS (coupling an UltiMate 3000 Nano LC system (Thermo Scientiic, Bremen, Germany) and an HTC-PAL autosampler (CTC Analytics, Zwingen, Switzerland) to a Q exactive plus mass spectrometer (Thermo Scientific)). Peptides were delivered to an analytical column (75 μm × 30 cm, packed in-house with ReproSil-Pur C18-AQ, 1.9 μm resin, Dr. Maisch, Ammerbuch, Germany) and separated at a flow rate of 280 nL/min using a 145-min gradient from 5% to 30% of solvent B (solvent A, 0.1% FA and 2% acetonitrile; solvent B, 0.1% FA and 90% acetonitrile). The instrument was operated in the data-dependent mode. Survey full scan MS spectra (m/z 350 to 1800) were acquired in the Orbitrap with 70,000 resolution after accumulation of ions to a 1 × 106 target value. Dynamic exclusion was set to 30s. The 12 most intense multiplied charged ions (z ≥ 2) were sequentially accumulated to a 1 × 105 target value and fragmented in the collision cell by CID with a maximum injection time of 120 ms. Typical mass spectrometric conditions were as follows: spray voltage, 2 kV; heated capillary temperature, 200 °C; normalized HCD collision energy, 25%. The MS/MS ion selection threshold was set to 2.5 × 104 counts. A 2.0 Da isolation width was chosen. Raw MS data were processed by MaxQuant (version 1.6.3.3) ^126^ supported by the Andromeda search engine. The MS/MS spectra were searched against the UniProt mouse database with the following search parameters: full tryptic specificity, up to two missed cleavage sites, carbamidomethylation of cysteine residues set as a fixed modification, and N-terminal protein acetylation and methionine oxidation as variable modifications. The false discovery rate (FDR) of protein groups, PSM were less than 0.01.

### RNA-seq & LET-seq Analysis

Gonocyte RNA-seq data was obtained from the Gene Expression Omnibus (GEO) under accession number GSE235429 ^27^. Reads were aligned to mouse genome (mm10) using Hisat2 with paired-end mode and --dta option ^127^ after being trimmed to 90bp using SeqKit ^128^. Reads aligned to genomic blacklist regions (mm10-blacklist.v2) ^129^ were removed and read counts on gene exons were calculated using featureCounts with -p -t exon -g gene_id options ^130^. Transcripts per million (TPM) were calculated for each gene to measure expression level. Differentially expressed genes (DEGs) were determined by comparing replicates of two stages using DESeq2 ^131^ after adding one to raw read counts to eliminate outliers. 21846 protein-coding genes on autosomes and sex chromosomes were identified according to Mus_musculus.GRCm38.102.gtf from Ensemble (https://asia.ensembl.org/Mus_musculus/Info/Index) for quantitative analysis. RNA-seq data of *Meiosin* KO and WT germ cells was obtained from DDBJ Sequence Read Archive (DRA) under accession number DRA008525 ^37^, and RNA-seq data of adult testis was acquired from the Gene Expression Omnibus (GEO) under accession number GSE55060 ^132^. RNA-seq data of *Ezh2* cKO and WT male PGCs was obtained from the Gene Expression Omnibus (GEO) under accession number GSE141182 ^63^. All data were processed according to the pipelines described in the corresponding reports. LET-seq data of oocyte and preimplantation embryo were downloaded from the Gene Expression Omnibus (GEO) under accession number GSE235547 ^107^, raw data were processed according to the pipeline described in the report. Reads aligned to exons instead of genes were calculated by featureCounts ^130^, TPM rather than spike-in normalization was applied to compare expression level with gonocyte RNA-seq data.

### scRNA-seq Analysis

The processed dataset of each gonocyte stage was imported to Scanpy for downstream analysis ^133^. One-third of the P3 dataset was randomly selected because of the excessive sample size, the resampled dataset was merged with others for subsequent analysis. Genes expressed in fewer than three cells as well as cells with fewer than 200 expressed genes were removed from the dataset. Further data filtering was performed to discard cells with the number of genes expressed in the count matrix fewer than 2000 or more than 8000, cells with the total counts exceeding 2000, as well as cells with the percentage of counts in mitochondrial genes below 1% or above 20%. Count depth was normalized to 1e4 per cell using “scanpy.pp.normalize_per_cell” function followed by log1p transformation using “scanpy.pp.log1p” function. Highly variable genes were defined using “scanpy.pp.highly_variable_genes” function with parameters “min_mean=0.0125, max_mean=3, min_disp=0.5”, and gene counts were scaled using “scanpy.pp.scale” with parameter “max_value=10” after mitochondrial genes were removed. Dimension reduction was conducted based on highly variable genes using “scanpy.tl.pca” function with parameters “use_highly_variable=True, svd_solver=’arpack’”. The nearest neighbors distance matrix was calculated using “scanpy.pp.neighbors” with parameters “n_neighbors=30, n_pcs=30” and dimensionality was reduced using Uniform Manifold Approximation and Projection (UMAP) by “scanpy.tl.umap” function. By employing the Leiden graph-clustering method using “scanpy.tl.leiden” function and annotating cell clusters with known marker genes, clusters of cell types other than germ cells were discarded. PCA dimension reduction for a filtered dataset containing only germ cells was conducted again with the same pipeline and gene expression in germ cells was demonstrated by performing sc.tl.umap with parameters min_dist=2, spread=5.

### Spike-in ChIP-seq Analysis

Reads were first trimmed to 67bp using SeqKit ^128^ and adaptors were removed using Cutadapt ^134^. Reads were then respectively aligned to the mouse genome (mm10) and exogenous DNA sequence provided by the manufacturer (EpiCypher, SNAP-ChIP® K-MetStat Panel) using BWA-MEM ^135^. For reads aligned to mm10, duplicates were dropped using Picard and those aligned to genomic blacklist regions (mm10-blacklist.v2) ^129^ were removed. Both reads mapped to exogenous or endogenous sequences were filtered with mapping quality > 30 using SAMtools ^136^. Spike-in scale factors were then calculated using ChIPSeqSpike ^137^ and bam files were converted to normalized bigwig files using deepTools function bamCoverage with --extendReads --scaleFactor options ^138^. Merged bam and bigwig files were obtained with the same pipeline for quantitative analysis. Bigwig files were also generated with both CPM normalization and spike-in normalization when compared with P7 undifferentiated spermatogonia H3K27me3 ChIP-seq data. H3K27me3 enrichment within promoters was defined by the normalized count of reads mapped to TSS +/- 500 regions using BEDtools function multicov ^139^.

### ChIP-seq & CUT&RUN Analysis

Gonocyte H3K4me3 ChIP-seq data at E13.5, E17.5 and E19.5, and H3K27me3 & H3K4me3 ChIP-seq data of P7 undifferentiated spermatogonia were downloaded from Gene Expression Omnibus (GEO) under accession no. GSE121118 ^21^ and no. GSE89502 ^79^, respectively. Gonocyte H3K4me3 ChIP-seq data was processed according to the pipelines described in the previous reports. Reads of P7 undifferentiated spermatogonia H3K27me3 & H3K4me3 ChIP-seq data were trimmed to 30bp and aligned to mm10 genome using BWA-MEM ^135^ after removing adaptors. Reads aligned to blacklist regions (mm10-blacklist.v2) ^129^ were removed and PCR duplicates were dropped with Picard. Bigwig files were obtained using deepTools function bamCoverage ^138^ with parameters --binSize 50 --extendReads 50 --normalizeUsing CPM and merged files were also generated for subsequent analysis. EZH2 & RNF2 CUT&RUN data was trimmed to 50bp and aligned to the mouse (mm10) genome using Bowtie2 with the -N 0 option ^140^, and only unique reads were used for downstream analysis. Reads aligned to blacklist regions (mm10-blacklist.v2) ^129^ and PCR duplicated were removed. Merged files were generated and converted to Bigwig files using deepTools function bamCoverage ^138^. H3K27me3 CUT&RUN data of PRC1 cKO/Control adult undifferentiated spermatogonia were downloaded from Gene Expression Omnibus (GEO) under accession no. GSE221944 ^12^ and processed according to the pipelines described in the previous report. Read counts within promoters (TSS +/- 500) of each processed data were calculated using BEDtools function multicov ^139^ and used for further analysis.

### DNA Methylation Analysis

BS-seq data was obtained from DDBJ under accession numbers DRA000607 ^6^ and DRA002477 ^22^. Raw reads trimmed to 50 bp were aligned to the mouse (mm10) genome and the methylation levels were called using BSseeker2 ^141^ with default parameters. CGmaptools ^142^ was used to calculate CG methylation levels within promoters and genome. GC% within promoters +/- 500bp around TSS was calculated using BEDtools ^139^ function nuc.

### Peak & PTG Definition

A customized peak-calling strategy specialized for broad histone modification was employed for H3K27me3 spike-in ChIP-seq and H2AK119ub CATCH-seq data. Narrow peaks were first roughly called using macs2 ^143^ for H3K27me3 and macs3 ^143^ for H2AK119ub with --broad –nomodel –nolambda parameters, then neighbor peaks with distance smaller than 2kb for H3K27me3 and 50kb for H2AK119ub were merged to broad peaks using BEDtools ^139^. Reads in peaks (RiP) were calculated and normalized with spike-in scale factors, then peaks with lower normalized RiP were filtered out to extract highly enriched sites. PTGs were defined by intersecting promoters of protein-coding genes with filtered peaks using BEDtools ^139^ with -u -f 1 parameters. Peaks for EZH2 and RNF2 CUT&RUN, as well as both peaks and PTGs for H3K27me3 ChIP-seq in P7 undifferentiated spermatogonia, were defined using a similar strategy using different specific thresholds and without spike-in normalization. Peak annotation was conducted using Chipseeker^144,145^.

## Data Availability

All data in this study have been deposited in the Gene Expression Omnibus (GEO) under accession number GSE272748 (Spike-in ChIP-seq), GSE272749 (CATCH-Seq), GSE272750 (CUT&RUN), GSE272751 (scRNA-seq). The MS raw data of chromatin proteome generated in this study have been deposited in the ProteomeXchange Consortium (https://www.proteomexchange.org/) via the Jpost, partner repository under accession ID PXD056741/<u>JPST003418</u> (https://repository.jpostdb.org/preview/1103519356709b9acdf992).

## Acknowledgments

We thank Drs. Martin Loza, Sung-Joon Park, Kenta Nakai, Shosei Yoshida, and all members of the Siomi laboratory for discussions and comments on this study. We thank Johji Nomoto and Shuji Ohshima for maintenance of the mouse strain. We also thank the One-Stop Sharing Facility Center for Future Drug Discoveries (The University of Tokyo) for FACS. This study was supported by Japan Agency for Medical Research and Development (AMED) Grant Numbers 21bm0704041h0003 (to S.Y.) and 22jm0210084h0003 (to S.Y.), MEXT KAKENHI Grant Number JP19H05466 (to M.C.S.), JSPS KAKENHI Grant Number 19K06616 (to S.Y.), 19H05754 (to A.I.), JP22H04925 (PAGS), the Takeda Science Foundation (to S.Y.), the NOVARTIS Foundation (Japan) for the Promotion of Science (to S.Y.), and the Astellas Foundation for Research on Metabolic Disorders (to S.Y.).

## Author Contributions

T.F., H.N., Y.U., A.I., and S.Y. conducted biochemical analyses using gonocytes. P.L. performed almost all bioinformatic analysis with some help from T.F., except for some initial analysis done by H.N., Y.U., and S.Y.. M.H. and S.N. conducted and supervised the analysis for SSC. M.S. and Y.S. conducted and supervised the scRNA-seq analysis. R.H. and J.A. conducted and supervised chromatin proteome analysis. S.S.K, J.R.D and A.A. supervised the bioinformatic analysis. S.Y. and M.C.S. conceived the project and designed the experiments. S.Y. and M.C.S. wrote the manuscript with input from all authors.

## Supplemental Information

Document S1. Figures S1–S7

