## Supplemental Figures for "Priming Epigenetic Landscape at Gene Promoters through Transcriptional Activation in Mammalian Germ Cells"

### Figure S1. Related to Figure 1

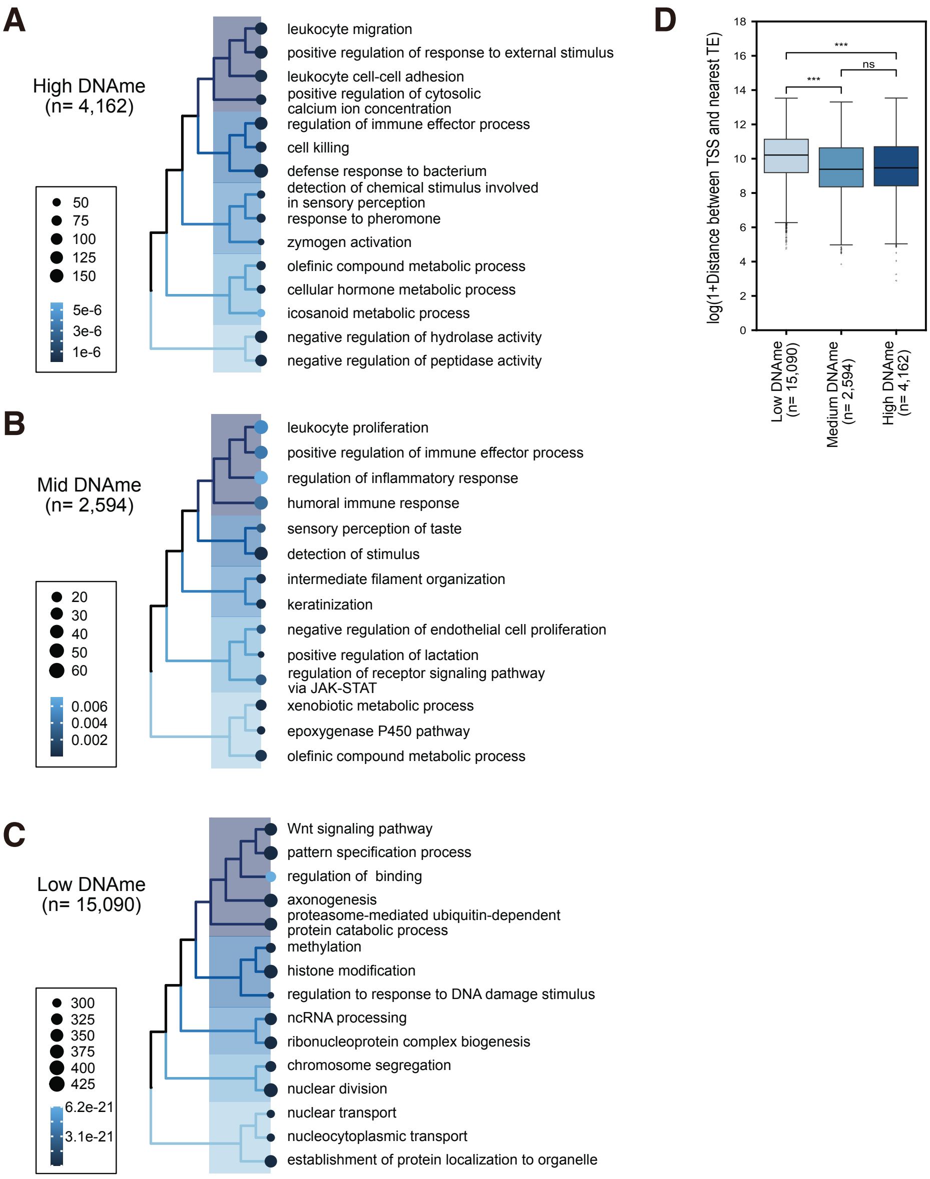


**Figure. S1: Profiles of Genes with Different Promoter DNA Methylation Levels in Gonocytes. Related to Figure 1.**

(**A**) Gene Ontology analysis with hierarchical clusters for High DNAme genes (n=4,162) defined at P0.

(**B**) Gene Ontology analysis with hierarchical clusters for Medium DNAme genes (n=2,594) defined at P0.

(**C**) Gene Ontology analysis with hierarchical clusters for Low DNAme genes (n=15,090) defined at P0.

(**D**) Distances between TSSs of genes and TEs on the genome of High, Medium, and Low DNAme genes defined at P0, respectively.

### Figure S2. Related to Figure 2, 3

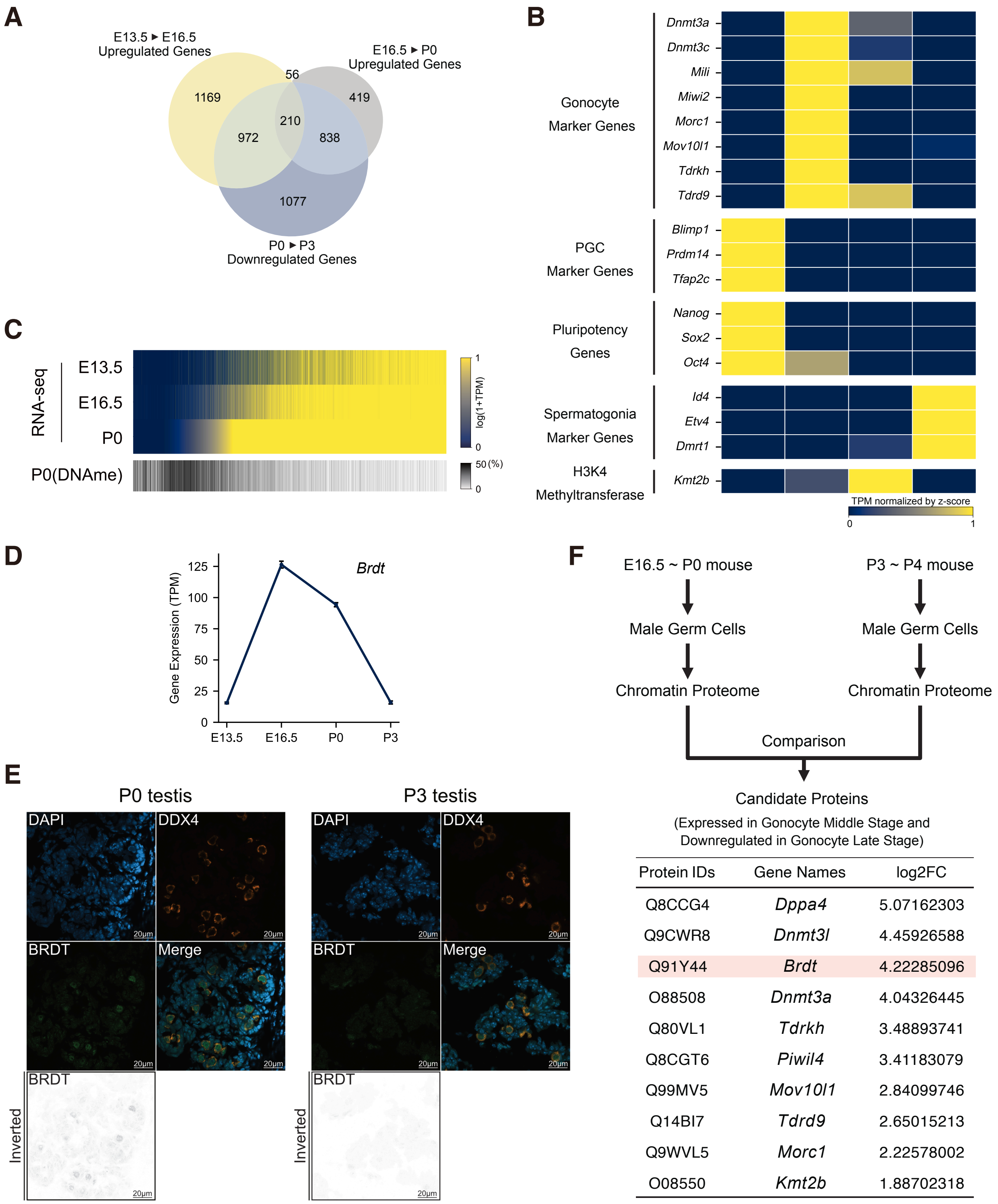


| Protein IDs | Gene Names | log2FC |
|---|---|---|
| Q8CCG4 | *Dppa4* | 5.07162303 |
| Q9CWR8 | *Dnmt3l* | 4.45926588 |
| Q91Y44 | *Brdt* | 4.22285096 |
| O88508 | *Dnmt3a* | 4.04326445 |
| Q80VL1 | *Tdrkh* | 3.48893741 |
| Q8CGT6 | *Piwil4* | 3.41183079 |
| Q99MV5 | *Mov10l1* | 2.84099746 |
| Q14BI7 | *Tdrd9* | 2.65015213 |
| Q9WVL5 | *Morc1* | 2.22578002 |
| O08550 | *Kmt2b* | 1.88702318 |

**Figure. S2: Transient and Ectopic Upregulation of Genes at Transcript and Protein Level. Related to Figure 2, 3.**

(**A**) Venn diagram illustrating the overlap between upregulated DEGs from E13.5 to E16.5 (n=2,407), upregulated DEGs from E16.5 to P0 (n=1,523) and downregulated DEGs from P0 to P3 (n=3,097).

(**B**) Heatmap showing gene expression levels of marker genes at different gonocyte developmental stages.

(**C**) Heatmap showing the correlation between gene expression levels at different developmental stages and P0 DNA methylation level, all genes are sorted by P0 gene expression level.

(**D**) Line plot showing transient gene activation of *Brdt* during the middle stage of gonocyte.

(**E**) Seminiferous tubule sections from P0 and P3 testis are stained for DAPI, DDX4 and BRDT, indicating the ectopic expression of BRDT protein during gonocyte middle stage.

(**F**) Schematic illustrating the pipeline of chromatin proteomics experiment and analysis in gonocyte middle and late stage, highlighting significant candidate proteins that are expressed in the middle stage and downregulated thereafter.

### Figure S3. Related to Figure 3, 4

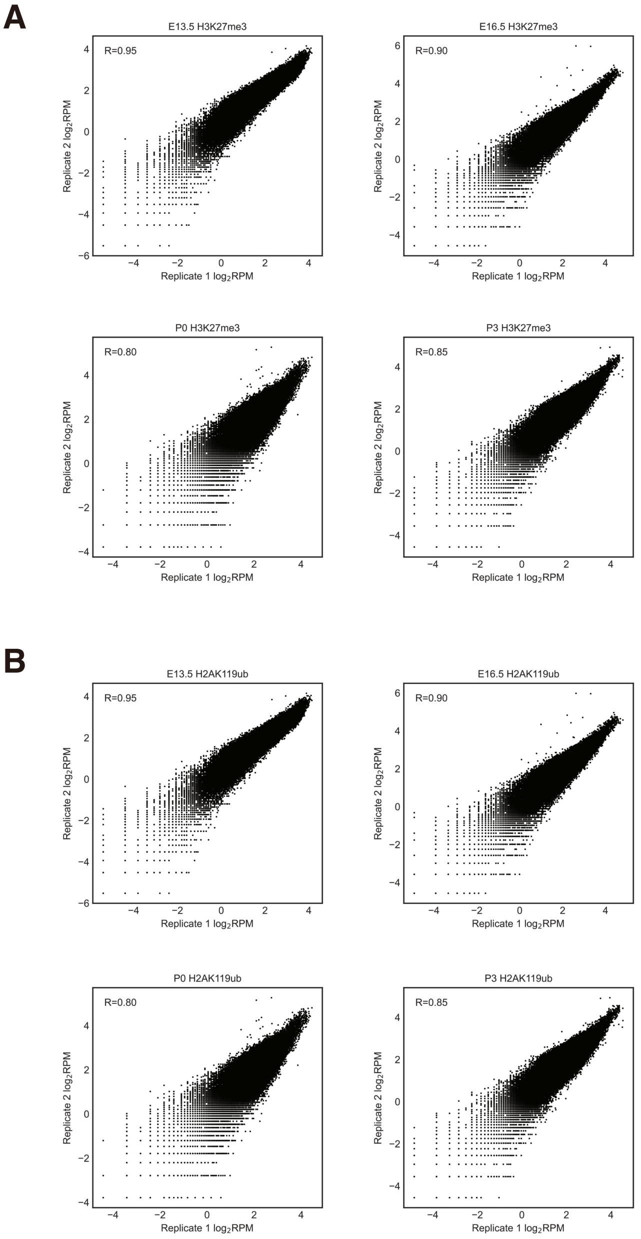


**Figure. S3: H3K27me3 Spike-in ChIP-seq and H2AK119ub CATCH-seq Dataset Analysis. Related to Figure 3, 4.**

(**A**) Scatter plots showing the correlation between biological replicates of H3K27me3 Spike-in ChIP-seq datasets at each stage, indicated by Spearman correlation of H3K27me3 enrichment across 10 kb windows.

(**B**) Scatter plots showing the correlation between biological replicates of H2AK119ub CATCH-seq datasets at each stage, indicated by Spearman correlation of H2AK119ub enrichment across 10 kb windows.

### Figure S4. Related to Figure 4

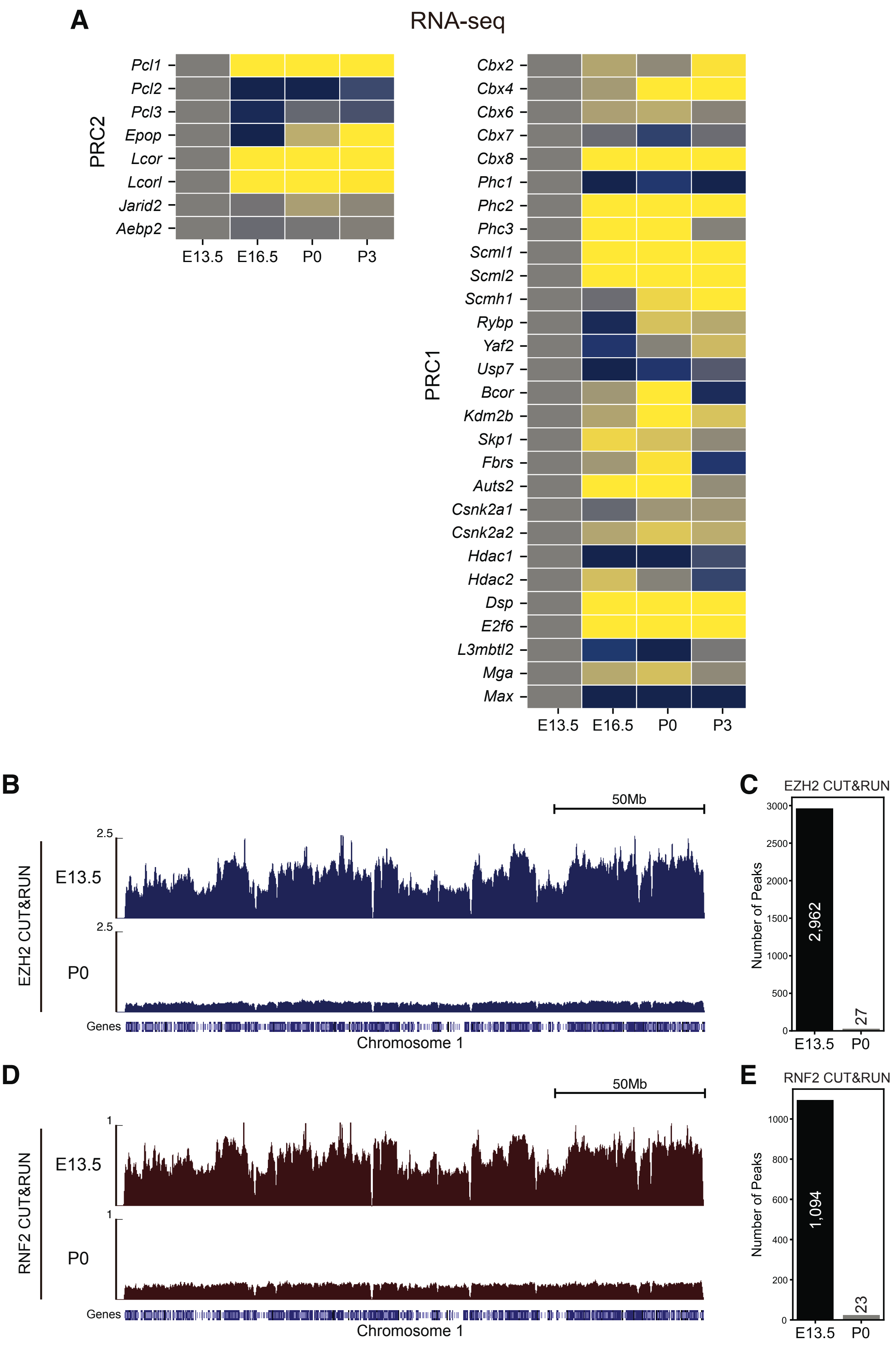

**Figure. S4: Gene Expression and Genomic Binding Analysis for PRC1/2 Components. Related to Figure 4.**

(**A**) Heatmap showing log2FoldChanges between gene expressions of each stage compared with E13.5, component genes of PRC2 and PRC1 are represented.

(**B**) Genome browser representations of Chromosome 1 showing EZH2 CUT&RUN merged datasets at E13.5 and P0.

(**C**) Number of peaks identified by EZH2 CUT&RUN at E13.5 and P0.

(**D**) Genome browser representations of Chromosome 1 showing RNF2 CUT&RUN merged datasets at E13.5 and P0.

(**E**) Number of peaks identified by RNF2 CUT&RUN at E13.5 and P0.

### Figure S5. Related to Figure 5

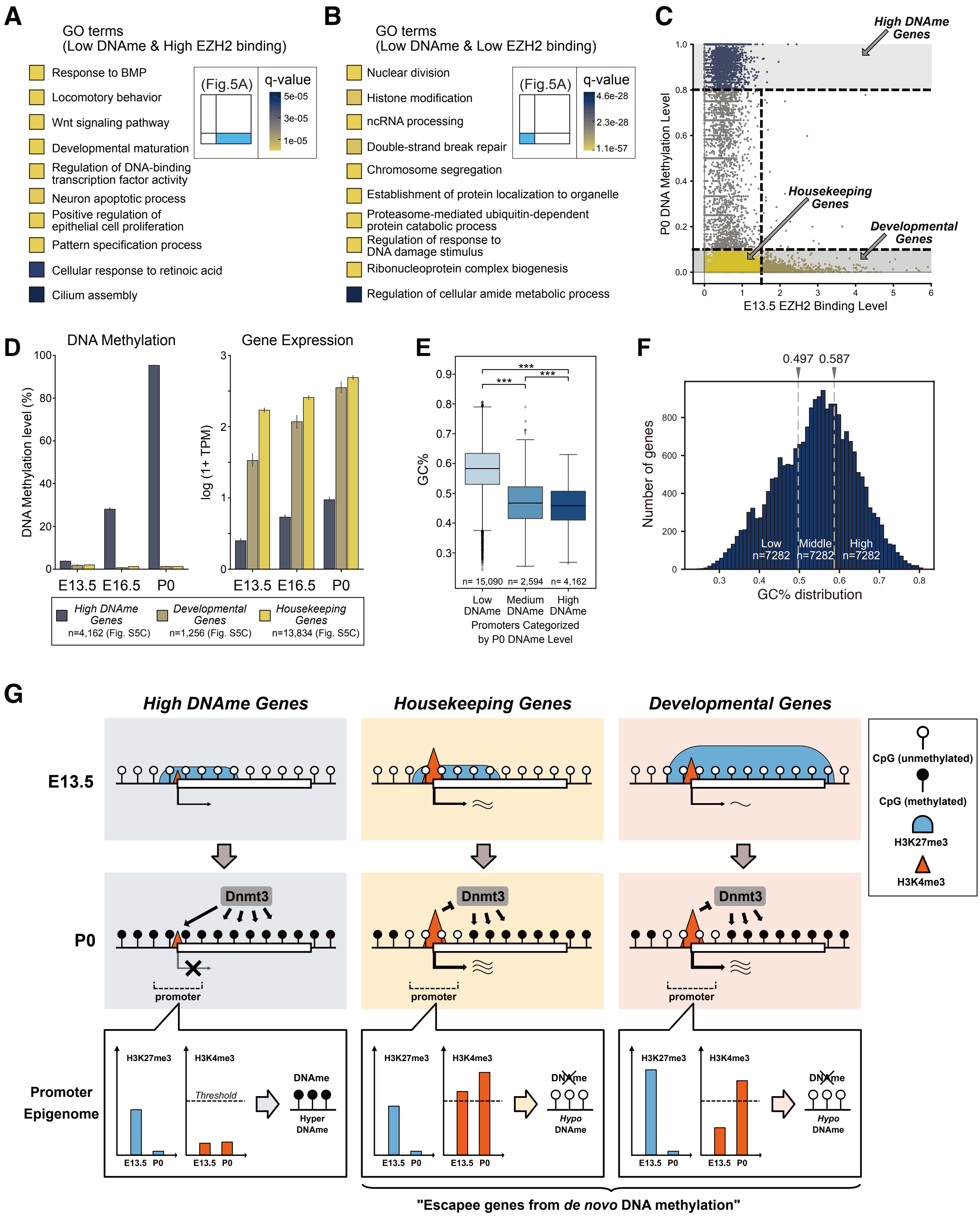

**Figure. S5: Combination of GC% and H3K4me3 Density Predicts *De Novo* DNA Methylation Levels at Promoters of Three Gene Groups. Related to Figure 5.**

(**A**) Gene Ontology analysis of genes with Low P0 DNAme level and High E13.5 EZH2 genomic binding level.

(**B**) Gene Ontology analysis of genes with Low P0 DNAme level and Low E13.5 EZH2 genomic binding level.

(**C**) Three groups of genes, namely *High DNAme Genes*, *Housekeeping Genes*, and *Developmental Genes* are defined based on P0 DNA methylation level and E13.5 EZH2 genomic binding level.

(**D**) DNA methylation levels and gene expression levels at each stage of three gene groups defined in (**C**).

(**E**) Boxplots showing GC% of genes in Low (0% - 10%), Medium (10% - 80%), and High (80% - 100%) DNAme groups at P0.

(**F**) Distribution of promoter GC% for all protein-coding genes, with thresholds dividing genes into three groups. The number of genes in each group is also annotated.

(**G**) Putative model illustrating how resetting histone modifications may contribute to shaping distinct epigenetic landscapes in gonocytes. During global reduction of repressive histone marks, *Housekeeping Genes* and *Developmental Genes* become highly activated, accompanied by increased H3K4me3 density antagonizing *de novo* DNA methylation. While *High DNAme Genes* experience no significant increase in gene activity or H3K4me3 density, eventually silenced by *de novo* DNA methylation.

### Figure S6. Related to Figure 6

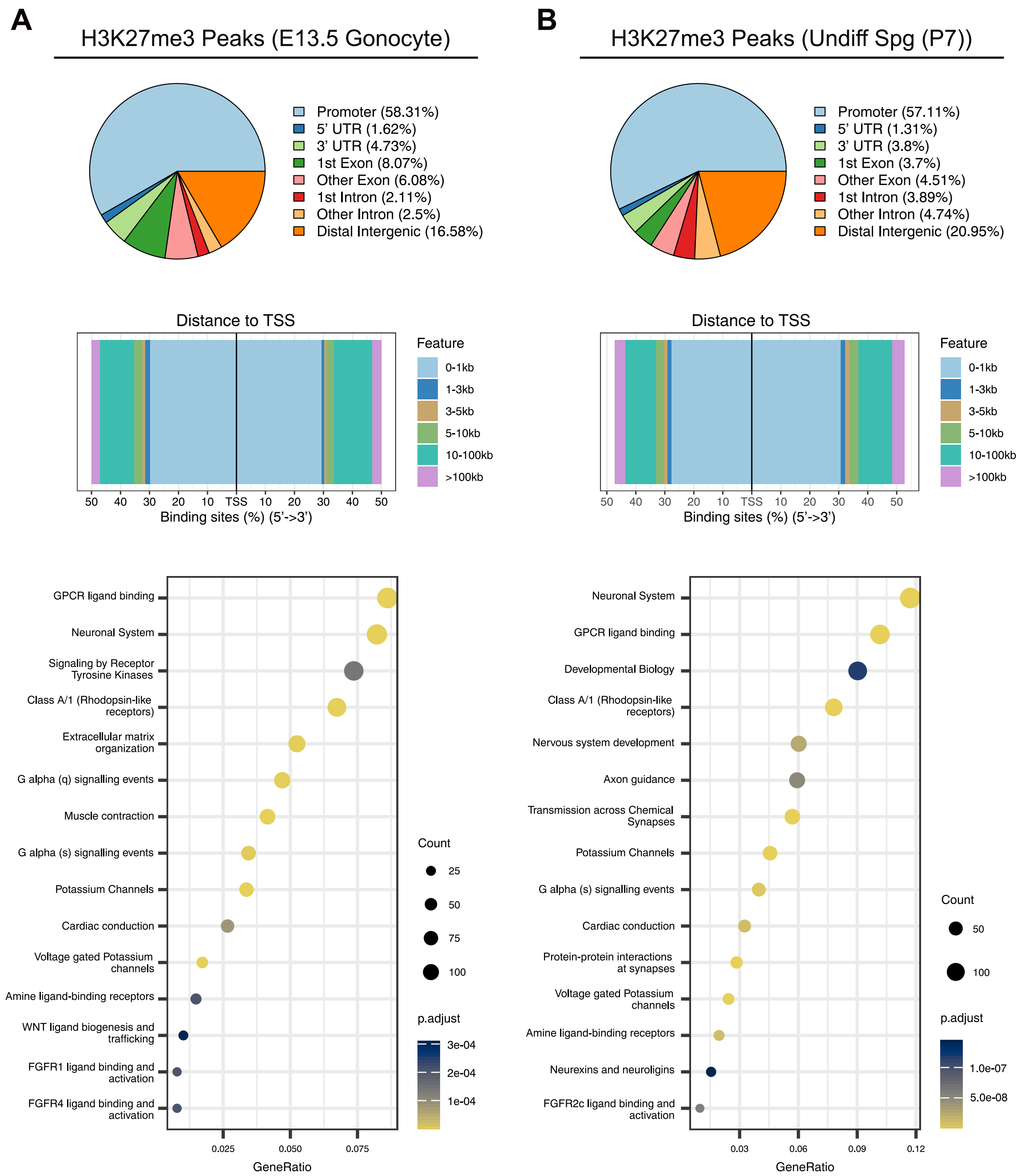


**Figure. S6: Comparison of H3K27me3 Peaks in E13.5 Gonocytes and P7 Undifferentiated Spermatogonia. Related to Figure 6.**

(**A**) Profiling the H3K27me3 peaks in E13.5 gonocytes, illustrating the distribution of peaks across different genome regions, their relative position to the TSSs, and the GO analysis of nearby genes.

(**B**) Profiling the H3K27me3 peaks in P7 undifferentiated spermatogonia, illustrating the distribution of peaks across different genome regions, their relative position to the TSSs, and the GO analysis of nearby genes.

### Figure S7. Related to Figure 7

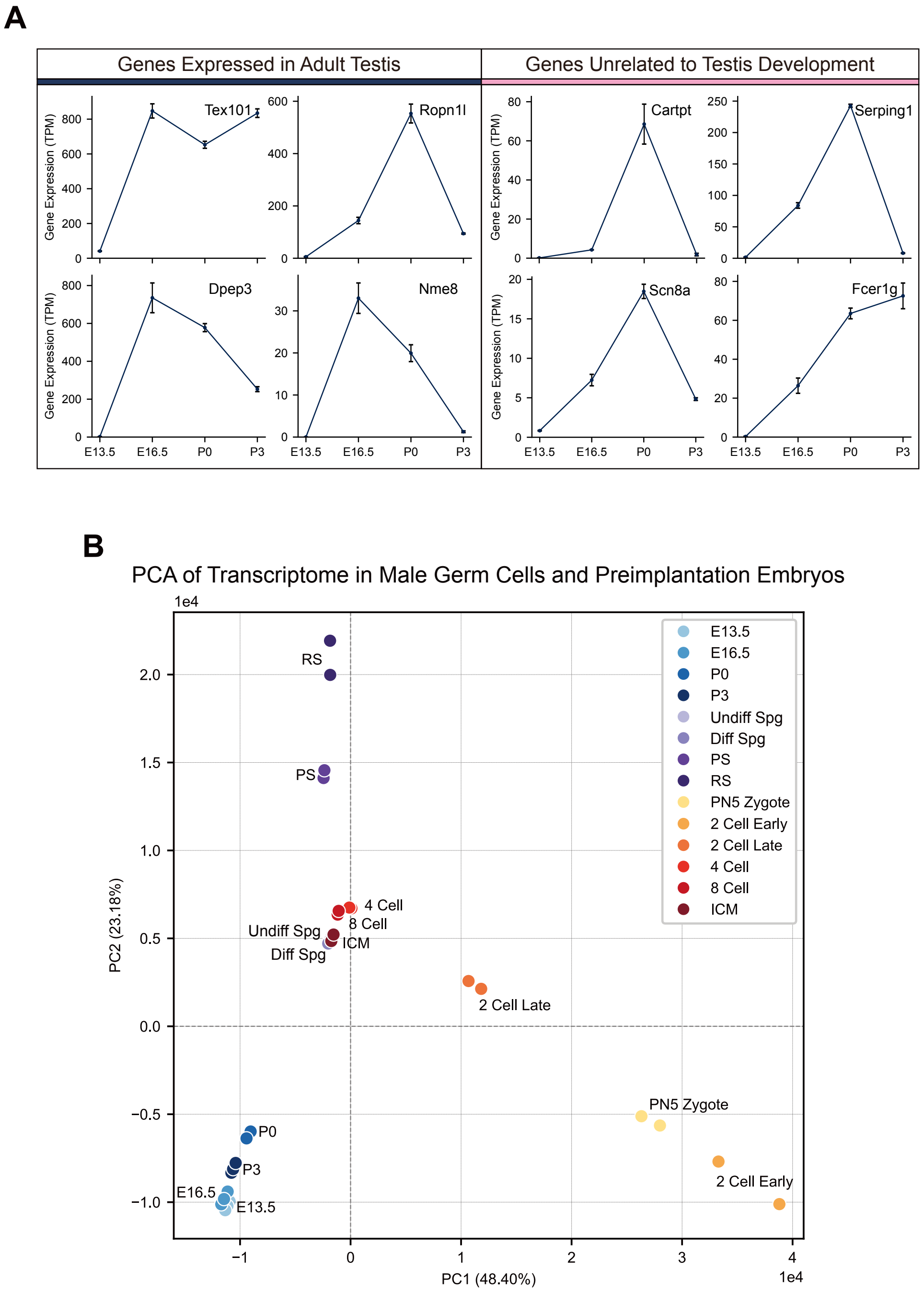


**Figure. S7: Transcriptome Analysis Reveals Ectopic Expression in Gonocytes. Related to Figure 7.**

(**A**) Gene expression levels at different gonocyte developmental stages analyzed using bulk RNA-seq data. Genes showing ectopic expression in gonocytes as shown in Fig. 7A are illustrated in two panels.

(**B**) PCA of transcriptome in male germ cells and preimplantation embryos. Undiff Spg: P7 undifferentiated spermatogonia, Diff Spg: P7 differentiating spermatogonia, PS: pachytene spermatocyte, RS: round spermatid, ICM: inner cell mass.
